# Towards a multi-stressor theory for coral reefs in a changing world

**DOI:** 10.1101/2022.03.22.485402

**Authors:** Carling Bieg, Henri Vallès, Alexander Tewfik, Brian E. Lapointe, Kevin S. McCann

## Abstract

Coral reefs are facing a constant barrage of human impacts, including eutrophication, overharvesting and climate change. While the local effects of overharvesting have been well-studied, regional nutrient loading from anthropogenic activities on land and global climate change-induced disturbances are increasing in magnitude and necessitating cross-scale multi-stressor approaches for coral reef ecology. Here, we expand on longstanding theory to develop an integrated multi-stressor framework for coral reefs. We show that: i) the geometry of a simple, empirically-motivated model suggests nutrients and harvesting can operate similarly, and synergistically, in driving shifts from coral- to algae-dominated reefs, resulting in clear context-dependent management implications; and ii) this same geometry suggests climate-driven coral mortality can drive the presence of long transients and climate-driven alternate states, even in moderately-impacted ecosystems. Reefs seemingly in a “safe space” based on individual stressors may in fact be much more susceptible to increasingly frequent storms and bleaching events in multi-stressor conditions. By integrating these findings with general ecological and theoretical concepts, we suggest that responses in benthic composition may act as “signatures of change” to multi-stressors, allowing us to develop a predictive and generalizable multi-stressor framework for coral reefs under global change. In line with this theory, we detail empirical evidence from Barbados of historical changes in reef composition and multi-stressor impacts within our framework. By bridging coral reef ecology and general ecological concepts, we can better understand ecosystem functioning and resilience in these important yet highly threatened systems.

**Manuscript Highlights:**

- Theoretical understanding of synergistic multi-stressor impacts on coral reefs
- Unexpected climate-driven alternate states, related to long transients
- Theoretical framework predicts “signatures of change” based on dominant stressor

## Introduction

Various axes of global change are increasing rapidly (Steffen and others 2015) and imposing a suite of multi-stressor impacts on ecosystems around the world, most notably coastal ecosystems such as coral reefs (Halpern and others 2019). For coral reefs, three major stressors have been identified as playing important roles in determining coral reef structure and ecosystem functioning: climate change, nutrient loading/water pollution, and overfishing, and these stressors have been studied extensively in isolation (Norström and others 2016; Harborne and others 2017; Hughes and others 2017). As coral reefs appear to be classic examples of ecosystems under heavy multi-stressor impacts (Scheffer and others 2015), it is important we deepen our understanding of the interactive effects of multi-stressors, across scales of space and time (Ban and others 2014; Pendleton and others 2016). For example, while climate change and temperature-induced coral bleaching is often, and understandably, targeted as the most important stressor threatening the future of coral reefs, local-scale stressors can significantly alter coral reefs’ resilience in the face of climate-related impacts (Wooldridge and Done 2012; Ban and others 2014; Abelson 2020; Donovan and others 2020). While addressing anthropogenic climate change at the global scale is undoubtedly critical, it is important we simultaneously consider the influence of regional-and local-scale management actions that may improve coral reefs’ resilience (Hughes and others 2017; Robinson and others 2019; Guan and others 2020; Donovan and others 2021). This is especially true if multiple stressors interact to nonlinearly increase negative impacts on coral reef function and would allow for a context-dependent approach to local management that could have dramatically more positive impacts.

At a local scale, the role of herbivory has been a dominant focus of coral reef research (Hughes and others 2007; Brandl and others 2019). Researchers have argued for decades that losses in herbivory can drive phase shifts from healthy coral-dominated reefs to degraded algal-dominated states (Mumby and others 2007), pointing to empirical examples of declining herbivores in reefs around the world to support this mechanism (Done 1992; Hughes 1994; Hughes and others 2007; Mumby and others 2007; Steneck and others 2014; Jouffray and others 2015). This has been argued to be caused by the overfishing of herbivores (McManus and others 2000; Hughes and others 2007; Mumby and Steneck 2008) as well as sudden reductions in other grazers not directly linked to fishing, such as the widespread *Diadema antillarium* die-off that occurred throughout the Caribbean in the 1980’s (Lessios 1988; Hughes 1994; Mumby and others 2007). Importantly, loss of hard corals and shifts to algae-dominated reefs result in highly degraded systems with the concomitant loss of important ecosystem services.

Adding complexity to the efficacy of local management actions aimed at restoring grazer populations, researchers and managers generally agree that coastal development and watershed pollution is a major local problem along with overfishing (Wear and Thurber 2015; Wear 2016). Nutrient loading has been shown to alter important biological parameters that mediate coral reef ecosystem dynamics, decrease coral calcification rates, and increase corals’ susceptibility to and severity of bleaching and disease (Bruno and others 2003; Wooldridge 2009; Wiedenmann and others 2013; Shantz and Burkepile 2014; Vega Thurber and others 2014; Zaneveld and others 2016; Wang and others 2018; Lapointe and others 2019; DeCarlo and others 2020; Donovan and others 2020; Houk and others 2022). Additionally, increased availability of nutrients have a positive impact on algae growth rates, and so nutrient loading can ultimately shift the dominant competitor from coral to algae (Lapointe 1999; Smith and others 2001).

Importantly, these collective biological responses to multiple stressors suggest there may be an interactive effect of nutrients, overfishing and climate variation in coral reef ecosystems and the competitive interaction between coral and algae (Burkepile and Hay 2006; Hughes and others 2017), and therefore context-dependent outcomes to management actions (Mumby and others 2006). Indeed, empirical research has found that the recovery of coral reefs after bleaching events appears to be highly dependent on multiple stressors including nutrient loading and herbivory (MacNeil and others 2019; Robinson and others 2019), and these stressors both appear to act together to change the rate and stages of algal succession following climatic disturbances (Hixon and Brostoff 1996; Mcclanahan 1997; Ceccarelli and others 2011). However, there still remains considerable debate among researchers regarding how multi-stressors interactively govern coral reef structure and functioning (Ban and others 2014; Muthukrishnan and Fong 2014; Cote and others 2016). Some researchers have begun to incorporate multiple stressors into theoretical and modelling approaches (Mumby and others 2006; Anthony and others 2011; Blackwood and others 2011, 2018; Fung and others 2011; Arias-González and others 2017), often done with either site-specific parameterizations and large amounts of detail or only incorporating one to two major stressors in more general ways. These studies have shown that multi-stressors can indeed interact and together contribute to increasing rates of hard coral decline, however disparity in approaches has prevented a generalizable framework for multi-stressor impacts on coral reef functioning.

While the nuances of coral reef dynamics and the multitude of anthropogenic impacts altering them are understandably not easy to discern, researchers have called for a holistic multi-stressor approach to coral reef science and management appropriate for a changing world (Norström and others 2009; Ban and others 2014; Pendleton and others 2016; Mumby 2017). Here, as a step towards this goal, we expand on existing theory (Mumby and others 2007) to demonstrate the context-dependency of management outcomes for coral reefs under multi-stressor impacts. We do this by integrating the combined influence of overfishing (i.e., loss/reduced herbivory), decreasing water quality (nutrient loading) and climate change (temperature stress and cyclonic storm damage) on coral reef structure and function within a theoretical multi-stressor framework (Figure 1A), and highlight the importance of highly responsive r-strategy life histories in changing, heavily impacted systems (Figure 1B). Our approach integrates general ecological concepts (e.g., ecological succession, keystone predation theory) with our current understanding of individual processes in coral reef ecosystem functioning and emerging theoretical insight into the role of noise in transient dynamics and ecosystem resilience. We end by evaluating empirical evidence from Barbados within our framework that corroborates the importance of considering an integrated multi-stressor perspective for coral reef management.

**Figure 1.**
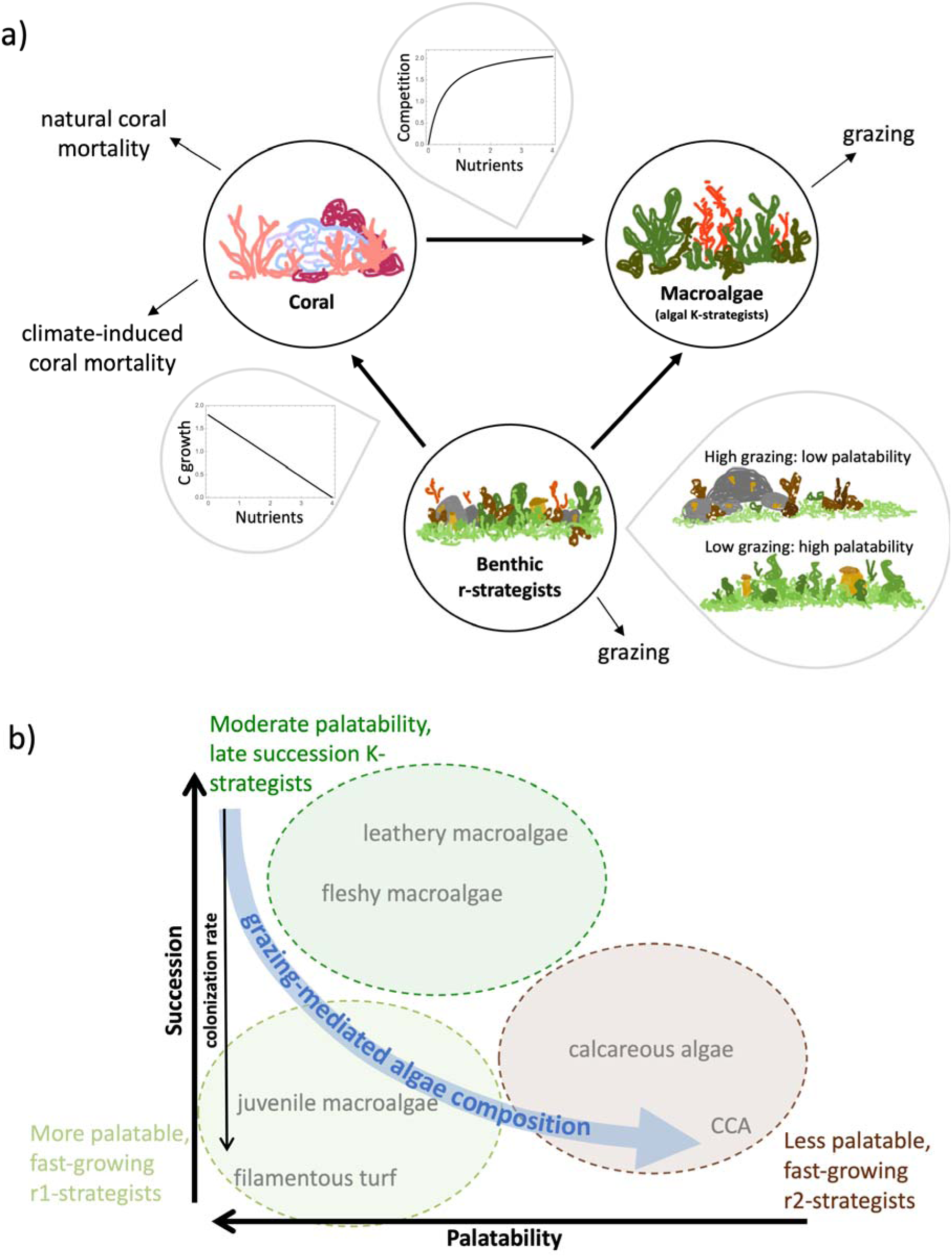
A) Summary of the model, interactions between state variables, and the incorporation of grazing (overfishing), nutrients and climate-induced mortality into scaling the model parameters. Increasing nutrients inhibit coral calcification (coral per capita rate of increase, *r,* decreases linearly with nutrients), as well as increase the growth rate of macroalgae and decrease coral’s competitive ability and resistance to macroalgae overgrowth (algal overgrowth rate, *a*, increases with nutrients up to a saturation point). Climate change alters coral mortality through stochastic events (repeated pulse perturbations at a defined frequency, akin to extreme climate events such as bleaching events (i.e., temperature stress) and destructive weather events like cyclonic storms). B) Schematic showing grazing-mediated algae composition in coral reefs based on growth-palatability trade-offs. High grazing rates suppress late-succession, palatable macroalgae while simultaneously selecting for less-palatable life strategies (through physical and chemical defenses) within early-succession algal r-strategists. Here we consider r-strategists as algae functional groups that colonize disturbed reefs quickly relative to thick, fleshy macroalgae (which are K-strategists relative to other functional forms of algae). We argue that within the group of benthic (B) r-strategists there are a variety of life history strategies that will be differentially selected for based on grazing rates. Note though that at moderate-high grazing rates, palatable fast-growing r-strategists may be able to withstand the grazing rates if their productive capacity simply outweighs grazing rates, however, at extremely high grazing rates (e.g., extreme *Diadema* densities), the relatively fast low-palatability organisms (e.g., CCA; r2) will thrive.

## Methods

Here, we synthesize multi-stressor components that have either been evaluated separately or in less general simulation approaches as an important step towards developing a general and mechanistic understanding of coral reef ecosystems under multi-stressor impacts. To do so, we expanded on a single-stressor (herbivory) model first introduced by Mumby et al. (2007) by incorporating the synergistic effects of overfishing (grazing), nutrient loading (algal competition and coral physiological stress) and climate change (coral mortality). This model has been widely used and extended by many researchers to including, but not limited to, additional complexity (reviewed in detail by Blackwood et al. (2018)), such as socio-ecological (Thampi and others 2018) and grazer dynamics (Blackwood and others 2011, 2012), additional benthic components (e.g., sponges) (González-Rivero and others 2011; Briggs and others 2018), spatial dynamics (Andréfouët and others 2002; Mumby 2006; Mumby and others 2006, 2014; Greiner and others 2022) and, like us, multiple stressors (Anthony and others 2011; Fung and others 2011; Arias-González and others 2017).

We follow recent evidence from the literature that suggests coral calcification is inhibited by increased nutrients (Shantz and Burkepile 2014), and similarly make the intuitive assumption that nutrients increase the growth rate of macroalgae (a primary producer) over coral (i.e., its growth rate), while simultaneously decreasing coral’s resistance to macroalgae overgrowth and competitive ability (Lapointe 1999; McClanahan and others 2003). These modifications to the original Mumby et al. model are discussed in more detail below. Figure 1A shows a schematic summarizing our model and the interactions between state variables. As such, our new nutrient-dependent model is as follows:

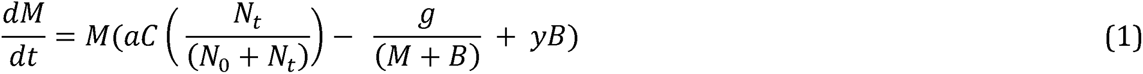

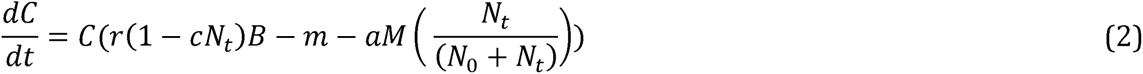

Here, M, C and B respectively represent macroalgae, coral, and benthic r-strategists that instantaneously fill empty space left by dead coral and grazed macroalgae. Since B is assumed to instantaneously fill any space (r-strategists act on much faster time scales relative to the other state variables), B = 1 – M – C and the model is therefore effectively reduced to two dimensions. Here, *a* is the rate that macroalgae overgrows coral, and thus can be thought of as a macroalgal-coral competition rate. *g* represents the grazing rate from herbivores, which is scaled by densities (or % cover since M + C + B = 1) of both M and B. *y* is the rate that macroalgae overgrows benthic r-strategists (B), *r* is the coral growth rate (again over B), and *m* is the natural mortality rate of coral.

### Nutrient effects

Nutrients (*N_t_*) alter the macroalgal-coral overgrowth rate (competition, *a*) in this model, based on the empirical evidence discussed above. We also assume that this nutrient-scaled competitive effect eventually saturates and therefore *N_0_* is a saturation constant that determines the effect of nutrients (*N_t_*) on macroalgal-coral competition (*a*). Additionally, we use the parameter *c* to represent the calcification rate – scaled by nutrient loading, *N_t_* – which therefore alters the coral growth rate (*r*) in response to nutrient loading. An important distinction between this model and Mumby et al.’s original model is that the parameter *a* can now be thought of as the maximum macroalgae-coral overgrowth rate (i.e., *a_max_*) and *r* similarly as the maximum coral growth rate (i.e., *r_max_*) due to their scaling by nutrients. That is, the realized growth rates depend on *N_t_* (Figure 1A).

While other studies have included nutrients in their models, this is generally done by only influencing algal growth rates (Mumby and others 2006; Anthony and others 2011; Hughes and others 2017). We note that since our model does not consider other aspects of water quality such as sedimentation, our results may be similar to others who have included this stressor in their models to negatively influence coral growth (Fung and others 2011; Gurney and others 2013; Arias-González and others 2017). However, sedimentation of course would have other effects on coral recruitment and various algal functional forms which we do not discuss here (Gurney and others 2013).

### Benthic r-strategists

The original model by Mumby et al. (2007) referred to this third variable as turf algae, a fast-growing r-strategist, however, we argue that nutrients and grazing rates would determine the composition of other components of benthic cover (Lapointe 1997; Lapointe and others 2018) (Figure 1A). Towards this end, we include another simple extension to represent algal life history traits. Specifically, we include a simple growth-palatability trade-off in algal community composition to explore the role of r-strategists in response to disturbances and grazing pressure (Figure 1). Here, we consider r-strategists as algal functional groups that colonize disturbed reefs quickly, relative to thick, fleshy macroalgae (late succession K-strategists relative to other functional forms of algae, but of course all are “fast” relative to corals), that will be differentially selected for based on grazing rates. High grazing rates favour less-palatable life strategies and fast growth rates that can withstand these grazing levels. Fast-growing r-strategists can withstand moderate-high grazing rates, explaining why we see a lot of turf algae and some less-palatable algae like CCA in some reefs. However, at extremely high grazing rates (e.g., extremely high *Diadema* densities), only low palatability organisms will be able to survive. While some unpalatable algal types like CCA are slow growing in terms of biomass, they are relatively quick colonizers and can thus be considered similar to r-strategists in the face of disturbances.

Thus, as a first approximation we include a linear 1:1 relationship between grazing rates and the proportion of more/less palatable r-strategists composing the benthic r-strategist guild. Here, we refer to highly palatable fast-growing r-strategists (e.g., filamentous turf) as r1-strategists, and slightly less palatable (but still fast-growing relative to K-strategists like fleshy macroalgae) r-strategists (e.g., CCA) as r2-strategists (Figure 1B).

Therefore, B = r1 + r2, where r2 = *g*B and r1 = (1-*g*)B.

These assumptions relate to well-known relationships between high grazing rates and less palatable algae (Sammarco 1982; Chiappone and others 2006), as well as research showing that grazing rates (among other drivers such as nutrients) can alter the rates and stages of succession in coral reefs (Hixon and Brostoff 1996; Mcclanahan 1997). Research has shown that algal community dynamics are important in overall coral reef ecosystem functioning (Renken and Mumby 2009; Renken and others 2010; Bozec and others 2016; Briggs and others 2018), and our goal is to integrate general ecological concepts into this literature (e.g., life history, keystone predation theory). Although a simple extension, our model formulation is a starting point for understanding the importance of algal life histories in general.

### Analysis

With Equations (1) and (2), we can symbolically solve for the model isoclines as follows:

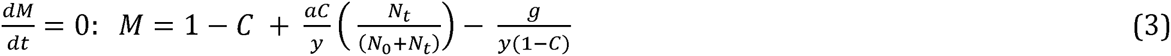

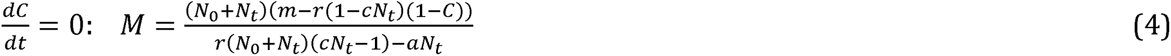

Immediately, we can see that both grazing (*g*) and nutrient loading (*N*) alter the isocline geometry (grazing via (3) and nutrients via both (3) and (4)) and thus equilibrium structure. These parameters both drive qualitatively similar changes in the isocline geometry (Supplement S.1) shown in Figure 2 and can cause both saddle node and transcritical bifurcations. The symbolic equilibrium solutions are not analytically useful, but this geometry is nonetheless useful in evaluating qualitative effects on equilibrium structure and bifurcations. Due to this geometric approach, the parameterization we chose was done to reflect the qualitative nature of possible outcomes in this model (i.e., transition through bistable parameter space) and explore the range of possible outcomes of coral reefs under global change. Any parameterization that gives this same qualitative geometry ought to display similar dynamics.

**Figure 2.**
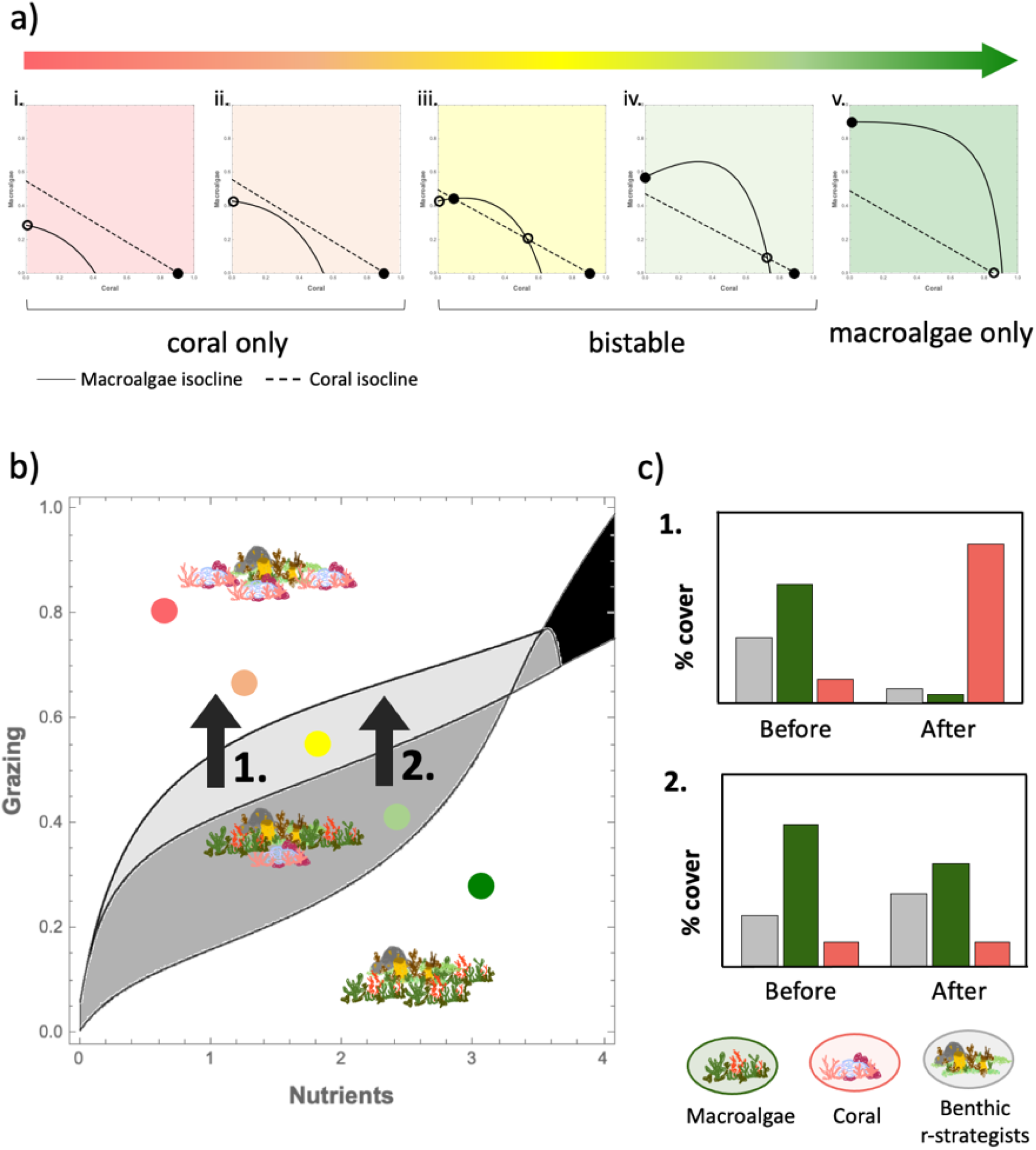
Interaction between grazing rates and nutrient loading on alternate states and management implications. A) Geometry of the model isoclines and equilibrium structure, and all possible (qualitative) configurations along a gradient of impact from coral-dominated reefs, through bistable configurations, to algae-dominated reefs. B) Regions of parameter space that correspond to geometrical configurations show in (A), highlighting regions of bistability. Grey regions show the area of parameter space where alternate states exist (yellow and light green), and the black region has only a 0-0 solution. Nutrients and grazing both drive a shift along this impact gradient, and they interact nonlinearly such that this shift is exacerbated. *a* = 2.0, *y* = 0.7, *m* = 0.15, *N_0_* = 0.5, *r* = 1.8, and *c* = 0.25. Black arrows (1 & 2) represent trajectories within this parameter space of management actions aimed at restoring herbivory and suggest context-dependent outcomes. C) Context-dependent restoration effects of management actions on benthic cover when reducing fishing pressure (e.g., MPA introduction) to increase grazing rates at low (1) and high (2) nutrient scenarios (represented by solid black arrows in B; 1, 2). Here, restoring grazers sufficiently ensures coral recovery in a low nutrient scenario (1) but remains in an algae-dominated state under higher nutrients due to hysteresis and failure to remove the algae-dominated equilibrium (2).

We incorporated climate effects (e.g., mortality events associated with cyclonic storm damage or mortality associated with bleaching events) using a flow-kick method (Meyer and others 2018), by inducing repeated pulse perturbations to coral cover (−50% cover at each occurrence) at fixed intervals, where the we were able to vary the frequency of disturbances to evaluate their effect on transients and asymptotic behaviour in comparison to deterministic simulations. All analyses were done using Wolfram Mathematica (version 12.1.0.0). Numerical simulations were evaluated using Mathematica’s built-in ODE solver “NDSolve” with integration method “StiffnessSwitching” (switches from explicit to implicit methods if stiffness is detected) when needed for non-deterministic simulations.

### Empirical Evidence

The Caribbean has a long history of research on coral reefs, and similarly long history of multi-stressor impacts, including widespread overfishing, catastrophic losses to urchin (important grazers) densities, eutrophication associated with agriculture and urbanization on land, and a history of other disturbances (Hughes and Connell 1999; Jackson and others 2014; Hughes and others 2017). Despite quantitative accounts of multi-stressor impacts being still somewhat hard to decipher, the Caribbean is an excellent starting point for integrating our theoretical framework developed here with empirical evidence. As such, we used historical data from reefs along the west coast of Barbados to evaluate temporal changes in benthic composition (coral and algal cover), grazer intensity (urchin density), and nutrient levels (various measurements), as discussed in further detail, including the natural and research histories of this area, below and in Supplement S.3.

## Results

### Towards a Multi-Stressor Theory for Coral Reef Ecology

Here, following empirical evidence, multi-stressors alter the biological rates and interactions included in the original model as illustrated in Figure 1A (also see Methods), and we explore the role of variations in faster algal life history strategies along a growth-palatability gradient (r1 and r2 compared to K-strategist, M) in response to multi-stressors (e.g., grazing resistance, growth rates), as demonstrated in Figure 1B. Towards understanding multi-stressors as an integrated whole, we start by first looking at the individual stressors of herbivory (top-down) and nutrients (bottom-up) in the absence of climate-driven coral mortality. We then examine the interaction of these three stressors together.

#### i. Bottom-up and Top-down: The Geometry of Nutrients and Grazing Impacts on Coral Collapse

Our choice of a simple model formulation allows us to use traditional phase plane techniques to analytically explore the dynamic outcomes of this coral reef model. As such, we can immediately see that both increasing nutrient loading and overfishing of herbivores (i.e., reduced grazing) drive a qualitatively similar sequence of dynamic outcomes in this model (Figure 2A). Assuming we start in a pristine coral reef scenario (pink coral zone; Figure 2A i), increasing nutrients or fishing pressure alter the geometry of the isoclines (see Supplement S.1 for individual stressor effects) driving bistabilty (yellow, light green, Figure 2A iii, iv) and finally complete loss of coral (dark green zone, Figure 2A v). This initial result suggests that bottom-up factors like nutrients alone, under an otherwise relatively pristine system (e.g., a marine protected area, MPA, or no-take zone), can shift coral reefs to an algae-dominated state (bistability, Figure 2A iii, iv), and with high enough nutrients drive the complete loss of the coral-only stable equilibrium (Figure 2A v). Thus, simple empirically-motivated extensions of this classic coral model find that both top-down and bottom-up impacts, in isolation, have qualitatively identical impacts on coral ecosystems.

This similarity in response immediately implies that when both stressors are increased simultaneously there is a strong tendency for the collapse of the coral state to occur under lower individual stressor – that is, overharvesting or eutrophication – values, compared to when each stressor is altered alone. Additionally, these stressors appear to have a synergistic (i.e., nonlinear) effect when in combination (Figure 2B; follow the colored dots to see a multi-stressor trajectory). While a simple geometric extension of Mumby et al. (2007), these results importantly suggest that we should expect context-dependent management outcomes, for example to the implementation of fisheries regulations (e.g., herbivore bans, no-take areas; Figure 2B,C), such that low nutrient cases may be more prone to success (e.g. Figure 2B, scenario 1) than nutrient enriched areas (e.g., Figure 2B, scenario 2). In other words, fishing regulations may be ineffective if other stressors, like nutrient levels, are too high (Figure 2C). We note this first result is aligned with Bruno et al.’s (2019) meta-analytic paper, which shows extremely variable coral reef ecosystem recovery responses to MPAs, strongly suggesting that other factors are involved and may even be the primary driver of coral reef structure and functioning (Suchley and Alvarez-Filip 2018), as well as other simulation results suggesting context-dependency of MPA success (Mumby and others 2006). This is not to say that herbivory is not important, however, an emphasis on it alone has little chance of significant success in the face of other stressors, and could lead to dangerous implications for ecosystem services provided by coral reefs and those whose livelihoods depend on their fisheries (Aronson and Precht 2006; Cline and Allgeier 2022).

#### ii. Long Transients and Climate-driven Coral Collapse

With our geometric understanding of how nutrients and fishing alter coral ecosystems we next turn to consider the impact of how climate intersects with these stressors to alter coral reef dynamics. We explore this by considering deterministic isocline arrangements that yield only a healthy coral reef state (i.e., the deterministic skeleton (Higgins and others 1997) is not bistable; Figure 3A,B i), but specifically focus on the very plausible scenario that there is some deterioration via both nutrients and fishing (i.e., Figure 2A ii.), which we will refer to as **moderate local impacts**. While under apparently “healthy” conditions (i.e., only a coral-dominated stable state), we note that the geometry implies that the two isoclines are relatively close to intersecting, and therefore near a saddle-node bifurcation (e.g., Figures 2A ii, 3A). Under such a biologically plausible condition (i.e, at least some human impact), we find that climate-induced coral mortality (i.e., adding stochasticity) greatly alters reef dynamics (e.g., Figure 3A,B ii).

**Figure 3.**
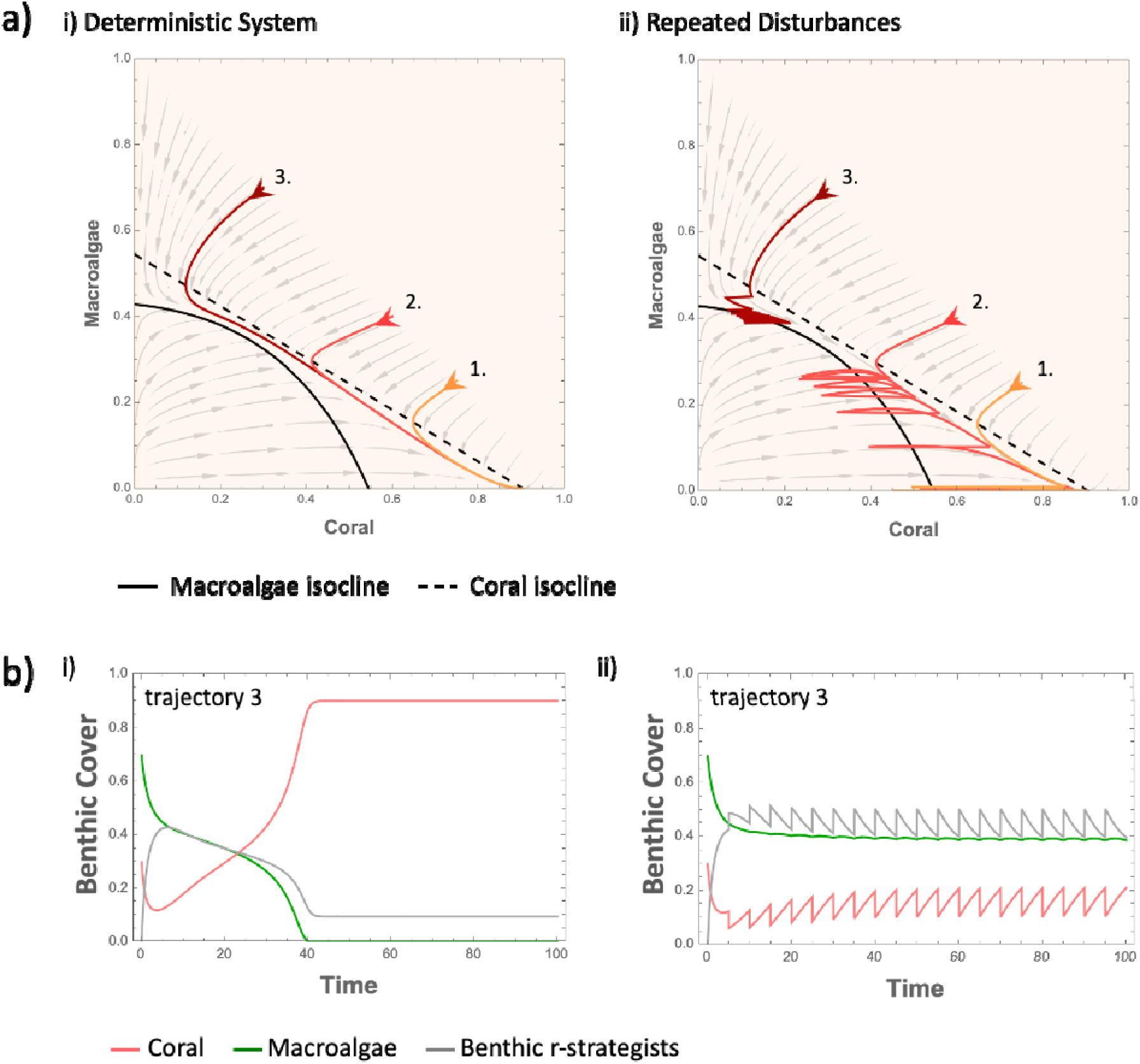
A) Isocline geometry and phase plane trajectories under moderate local impacts in (i) deterministic (no disturbances) and (ii) perturbed simulations (recurring climate-induced coral mortality events) starting at different initial values (numbered trajectories). Since there is only one stable equilibrium (the coral axial equilibrium), all deterministic solutions (i) eventually reach this equilibrium. However, we can have drastically different outcomes in the face of repeated climate-induced disturbances (ii). Note this state (moderate local impacts) is near a bifurcation point where nutrients and grazing can quickly drive a saddle-node bifurcation, creating two new equilibria (one stable and one unstable) where the isoclines intersect. Our trajectories appear be influenced by a nonexistent (but soon to be real) saddle point (the unstable equilibrium that appears after a saddle-node bifurcation), such that the vector field effectively has “memory” of it (notice the vector field – grey arrows – pulls toward this region). Behaviour like this has been referred to as a “ghost attractor” where the system mimics – or slows down – the dynamics as if the attractor were there (Hastings and others 2018; Morozov and others 2020). Here, we see that ghost attractors can cause long-transients and disturbance-driven alternate states. B) Time series of trajectory 3 (from part A i and ii) showing that benthic r-strategists flourish during these transients and periods of coral suppression. Here, *a* = 2.0, *y* = 0.7, *m* = 0.15, *g* = 0.4, *N_t_* = 0.53, *N_0_* = 0.5, *r* = 1.8, and *c* = 0.25. In ii) disturbance frequency is every 5 time units, where each perturbation results in an instantaneous loss of 50% coral cover.

First, even without climate-induced mortality events, we see that the time to reach equilibrium (the coral axial solution) is highly dependent on initial values (Figure 3A i), such that in certain cases (i.e., trajectory 3) the system is initially pulled to lower coral densities before eventually approaching the equilibrium. We also see that benthic r-strategists play an important role during these transient periods (Figure 3B i), something we elaborate on more below. Second, when adding climate-induced coral mortality events under moderate local impacts, not all trajectories have the same outcome (Figure 3A ii). Here, once again depending on the initial values, some trajectories get stochastically “entangled” in the region where the isoclines are nearly intersecting (dark red, trajectory 3), while others avoid this and eventually approach the coral axial equilibrium (though they notably take longer due to the perturbations). This result importantly suggests the potential for **climate-driven bistability**, such that there appears to be two different outcomes once climatic disturbances are incorporated into the simulations. Again, Figure 3B ii shows this third trajectory as a time series, where the asymptotic behaviour reflects an algae-dominated alternate state. Here, some coral persists but the system is largely dominated by macroalgae and benthic r-strategists. Note that while we focus here on a state of moderate impact, disturbances could of course cause local extinctions of coral in more degraded (i.e., bistable) ecosystems by knocking the system out of the basin of attraction to the coral equilibrium (bistable configurations in Figure 2A iii and iv; Supplement S.2).

Taken altogether, these results suggest that climate-driven coral mortality has the potential to greatly increase the presence of bistability, especially in a world already plagued by varying levels of nutrient loading and overfishing. Effectively, disturbances shift the resilience boundary for coral reefs (Meyer and others 2018), and the interaction of all three stressors operates nonlinearly to decrease coral reef resilience. Furthermore, increasing frequency of coral mortality events increases the region of state space where this climate-driven occur, meaning they may become more common under climate change (Supplement S.2). Note that there is a possibility that some of these stochastic trajectories, such as trajectory 3 in Figure 3, are extremely long transients that eventually squeeze through this region of entanglement (our climate-driven alternate state) if given enough time or variability in disturbance timing or severity. However, on ecological and management time scales this result remains functionally bistable. Depending on initial values and the frequency of disturbances, we can indeed see extremely long transients that eventually reach the coral-only attractor (Supplement S.2). On the other hand, coral collapse may have the potential to take an extremely long time due to the length and trajectories of these transients. If this is the case, it may seem like coral might persist, or even recover, before eventually disappearing (Supplement S.2), which would importantly suggest the possibility of extinction debt (Tilman and others 1994) in degraded coral reef ecosystems (i.e., whereby species are doomed to slow extinction even if the ecosystem deteriorates no more). This interesting result suggests that the interactions between moderate levels of multiple stressors may enhance the likelihood of bistability and long transients, both states that are incredibly hard to manage.

Interestingly, we can begin to explain these unexpected results using the geometry of our simple model (Figure 3A). While beyond the scope of this paper to fully investigate, we note that these results are consistent with phenomena recently described in the theoretical literature (Hastings and others 2018, 2021; Morozov and others 2020). Specifically, we see an interesting example of what has been referred to in the theoretical literature as a “ghost attractor”, such that the system is influenced by what is effectively the memory of a nonexistent equilibrium – in this case a saddle point (Figure 3). Hastings et al. (2018) and Morozov et al. (2020) provide more detailed information on long transients and ghost attractors, as well as saddle crawl-bys once the saddle node is present (after a bifurcation). Importantly, both of these phenomena can lead to long transients and slow shifts between states, which can have important implications in the face of stochasticity and we indeed see here (Figure 3). Notably, the geometry of this intermediate multi-stressor state explains both the change in direction of the trajectories shown in Figure 3A, as well as the “slow down” causing long transients and the emergence of a climate-driven algae dominated state seen in Figure 3A,B ii.

#### iii. Algal Trait Responses

Finally, to characterize different signatures of multi-stressors, we show how the relationship between stressors impacts algal community composition. Figure 3B shows that the role of benthic r-strategists (B) is important during long transients following a disturbance (Figure 3B i), as well as when frequent disturbances lead to a climate-driven alternate state (Figure 3B ii). Here, we once again focus on the case of moderate local impacts (Figure 2A ii), to examine how multiple stressors interact to influence the benthic algae composition in this state of uncertainty. Depending on grazing intensity, nutrient levels and frequency of climate perturbations, different benthic r-strategists (B) will dominate primary succession following large mortality events (Figure 4).

**Figure 4.**
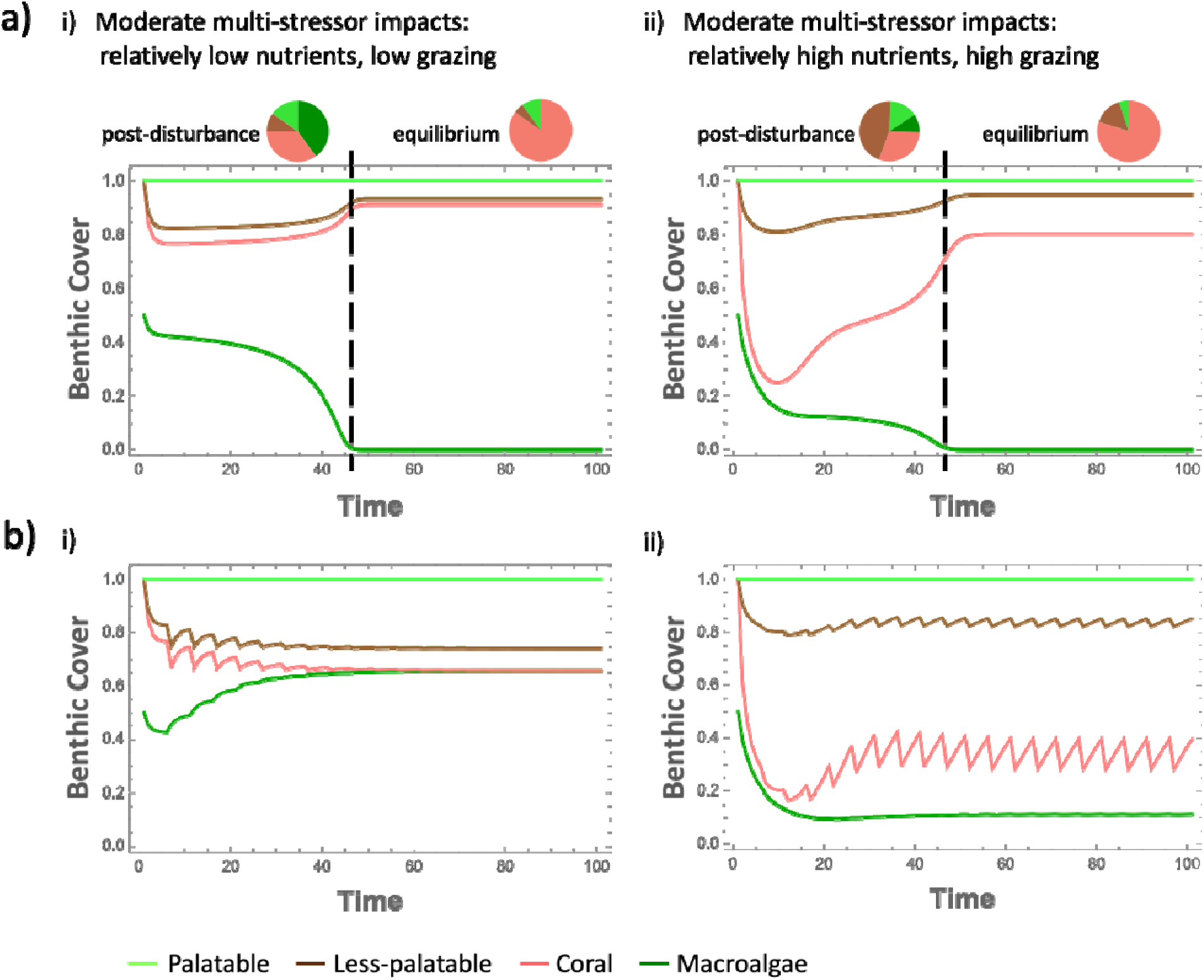
Benthic cover in a moderately impacted system (long transients after a perturbation) is altered by multiple stressors, through grazing-mediated algae composition and differential trajectories of coral recovery. Higher grazing rates select for fast-growing but relatively less palatable benthic r-strategists (e.g., r2 strategists such as CCA) that can productively withstand grazing, whereas lower grazing rates allow for more palatable (and likely even faster growing) benthic r-strategists dominate (e.g., r1 strategists like turf algae). Grazing of course also suppresses palatable macroalgae (K-strategist). A) The effect of single perturbations on benthic composition during transients (post-disturbance), and B) frequent climate-driven mortality events, such as temperature-induced bleaching events and cyclonic storm damage, can permanently alter benthic composition. Simulations in A) and B) have the same parameter values. In all cases *a* = 2.4, *y* = 0.7, *m* = 0.15, *N_0_* = 0.5, *r* = 1.8, and *c* = 0.25. Nutrients and grazing are varied as follows: i) *g* = 0.24, *N_t_* = 0.16; and ii) *g* = 0.7, *N_t_* = 2.31. In B) disturbance frequency is every 5 time units, and results in an instantaneous loss of 50% coral cover.

To elucidate these outcomes within our model, we performed a single climate pulse perturbation under different multi-stressor conditions (i.e., relatively low grazing and low nutrients versus relatively high grazing and high nutrients), within this geometrical configuration reflecting moderate local impact (Figure 4A). We see that grazing-mediated algae selection strongly influences the composition of benthic algae cover (r1-(i.e., highly-palatable, fast growing) vs. r2-(less-palatable, fast growing) vs. K- (palatable, slow growing) strategists), and all three stressors – nutrients, grazing, and climate – mediate the relative dominance of total benthic cover including coral (Figure 4A; see both post-disturbance transient responses as well as equilibrium composition). Notably, these results are exacerbated when frequent climate-induced mortality events are considered (Figure 4B). Frequent disturbances can permanently alter benthic composition by continuously suppressing coral recovery and impeding full algal community succession, while also compounding the effect of multi-stressors on the relative dominance of benthic composition (Figure 4B). Note that in Figure 4B i we see that frequent disturbances in a relatively low-nutrients, low-grazing scenario drive coral to extinction and cause a shift to a macroalgae-dominated state. In the relatively higher-nutrient, higher-grazing scenario shown in Figure 4B ii, coral can persist but is largely suppressed to the benefit of (mostly low-palatability) benthic r-strategists. These results again demonstrate the existence of climate-driven bistability seen in Figure 3, making this region one of high uncertainty given the interactions between multi-stressors. Importantly, our theory suggests that monitoring benthic cover responses ought to allow us to differentiate the role of multi-stressors and aid management decisions.

### Theoretical Synthesis: A framework for coral reef composition in a changing world

We have seen that multi-stressors have the potential to cause interactive, synergistic and non-linear effects, as well as potentially unexpected outcomes in a changing world. Our theoretical results also predict that moderate local impacts ought to generally display unexpected bistability with increasing climate variation. Indeed, increasing the frequency of climate-induced mortality events enlarges the area of bistability, therefore decreasing the resilience of coral-dominated reefs and increasing the chances of algal dominance (i.e., grey region in Figure 2B; see Supplement S.2 for simulation data). As a result, our region of moderate local impacts in parameter space (Figure 2) becomes a region of **climate-driven uncertainty**, which is likely characterized by a large presence of benthic r-strategists (r1 and r2) when faced with real-world climatic noise and perturbations. Here, the eventual fate of a system within this region of uncertainty will depend on a combination of disturbance frequency, severity, and history of the system itself. Additionally, climatic variation increases the likelihood of climate-induced state shifts (Supplement S.2), essentially meaning that all regions of bistability and algal dominance in the deterministic model will likely lead to algae-dominated reefs.

Here, we summarize our theoretical results as a multi-stressor framework for coral reefs in a changing world as an update to the nearly 40-year-old Relative Dominance Model (Littler and Littler 1984; Lapointe 1997), highlighting this important region of climate-driven uncertainty and noting the importance of multi-stressor impacts on determining benthic composition (Figure 5). Importantly, algal composition gives us a signature of the degree of top-down control in coral reefs. Specifically, high amounts of relatively less-palatable benthic organisms like CCA suggest significant top-down grazing pressure and therefore point to the fact that other stressors – such as climate and/or nutrient loading – are likely playing a larger role in driving the proliferation of algae and reduction of coral. In rapidly changing environments, this framework allows us to make qualitative predictions about the trajectories of benthic composition change – in other words, signatures of change – in coral reefs based on different levels of multi-stressor impacts.

**Figure 5.**
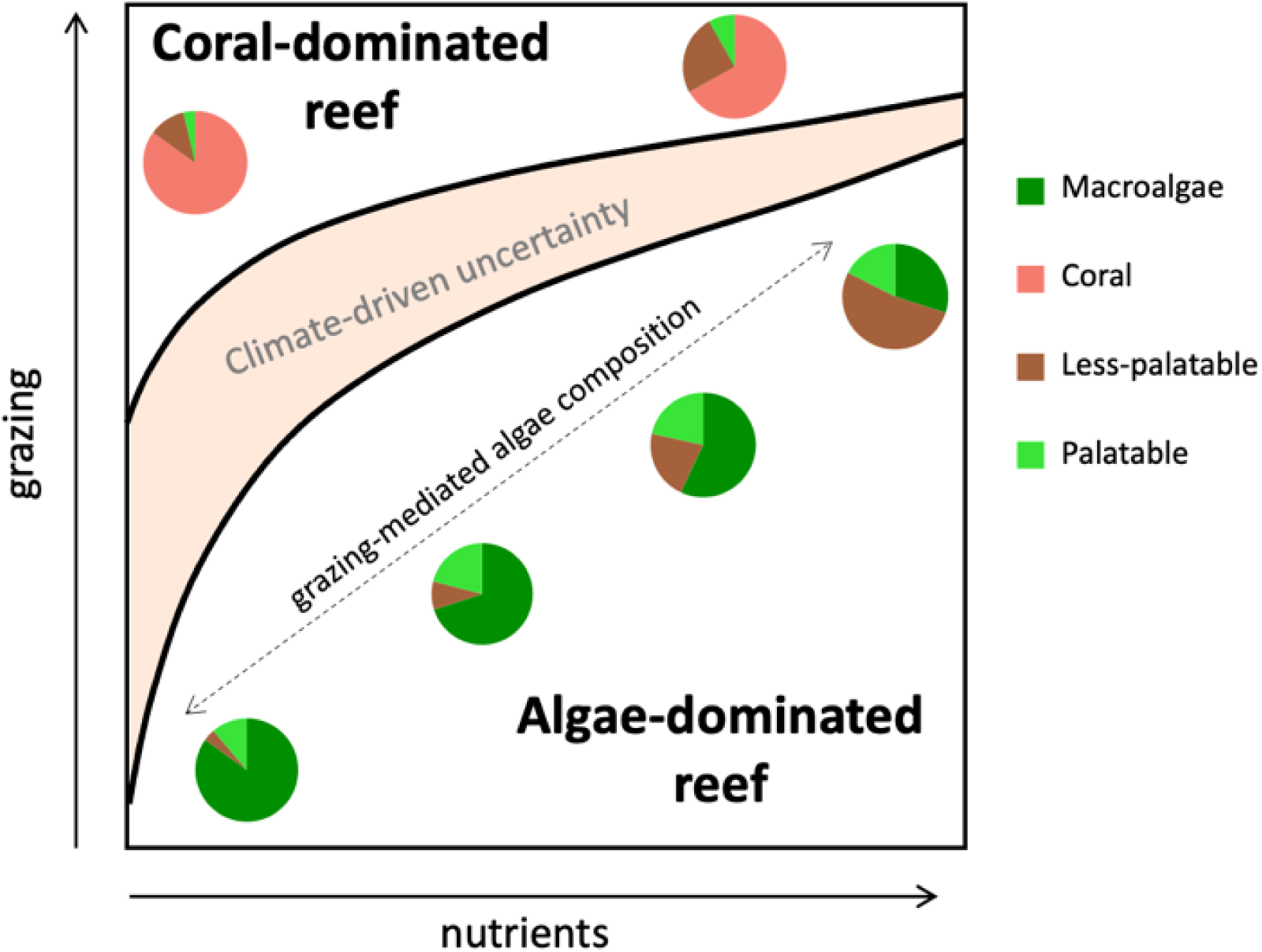
A summary of our theoretical results as a framework for coral reef composition in a changing world. Reef benthic cover depends on grazing, nutrients, and climate-related coral mortality. The region of multi-stressor space for coral-dominated reefs decreases with increased disturbance frequency (replaced by a region of climate-driven uncertainty), while the composition of algae (both slower-growing and late succession thick macroalgae, as well as early succession r-strategists) is mediated by grazing rates. Here, pie charts display the changing nature of benthic composition due to grazing-induced selection. Increasing disturbance frequency reduces the possibility of coral persistence by increasing the susceptibility to other stressors, decreasing the basin of attraction to the coral-dominated equilibrium when in a bistable configuration, and increasing the likelihood of climate-driven state shifts. We therefore argue that anything below the region of climate-driven uncertainty would be an algae-dominated system in a noisy world.

### Linking empirical evidence to our theoretical framework

Our theoretical framework importantly allows us to test predictions about changes in benthic cover under multi-stressor impacts and identify underlying drivers of change in coral reefs. As such, we now highlight evidence from the Caribbean within our theoretical framework. Reefs across the Caribbean notably underwent significant changes in the 1970’s and 1980’s with a general loss of hard corals (Gardner and others 2003) corresponding with the culmination of disease (e.g., white-band disease; Aronson and Precht, 2001), damaging hurricanes (e.g., Hurricane Allen in 1980), the widespread loss of *Diadema* in 1982/83, as well as ongoing eutrophication from untreated wastewater and agricultural runoff, and widespread overfishing in the region (Jackson and others 2014). As a first step of linking our theoretical framework to empirical accounts of changing reef conditions, we do this by integrating multiple studies from reefs along the west coast of Barbados (Supplement S.3) that demonstrate trends in nutrients and grazing rates and associated changes in benthic cover.

The west coast of Barbados has a long history of human impacts including heavy fishing and nutrient runoff from urban and agricultural areas (Allard 1994; Bell and Tomascik 1994; Tosic and others 2008; Gill and others 2019). Barbados also has a unique research history, in the sense that researchers have focused on both top-down (grazing) and bottom-up (nutrients) dynamics. Collectively, and with hindsight, this allows us a relatively broad perspective of the changing coral reefs during this period of drastic change.

Baseline accounts (qualitative descriptions (Lewis, John 1960) and quantitative surveys (Stearn and others 1977)) of the west coast’s fringing reefs suggest that live coral cover began declining as early as the 1960’s, with particularly drastic declines throughout the 1970’s, during which time algae began taking over (Tomascik and Sander 1987a) (Figure 6A,B). Coral was further depleted following Hurricane Allen in 1980 along the entire coast (Mah and Stearn 1986) (Figure 6B). At this time (1982/83), and prior to the widespread *Diadema* die-off, Tomascik and Sander began sampling multiple reefs along the west coast in an attempt to understand the major drivers of change in Barbados’ rapidly degrading coral reefs (Tomascik and Sander 1985, 1987a, 1987b) (Figure 6A; note high rates of change in Figure 6B). As a result of rapid development on land (Tomascik and Sander 1985) these surveys identified a north-south gradient of increasing eutrophication and decreasing *Diadema* densities, though average *Diadema* density was still quite high, even in the southern reefs (see Supplement S.3, Figure S8) (Tomascik and Sander 1987a). Along this eutrophication/grazing gradient, coral diversity, percent cover, growth, and overall recruitment all decreased (Tomascik and Sander 1987a, 1987b; Tomascik 1990, 1991; Hunte and Wittenberg 1992; Wittenberg and Hunte 1992; Mann 1994), however CCA cover – relative to turf and frondose macroalgae combined – increased with increasing *Diadema* densities (Supplementary Figure S8). These local patterns support our theoretical predictions, namely that coral was already rapidly declining before the reduction in grazing, possibly due to elevated nutrient levels, and that the still relatively high grazing rates meditated benthic algae composition locally. Note that despite the apparent gradient, the majority of the coastline may have been nutrient saturated by the early 1980’s (Tomascik and Sander 1985; Allard 1994), if not earlier (Sander and Moore 1979), based on various nutrient thresholds (Supplement S.3) (Lapointe 1997; Bell and others 2014; Lapointe and others 2019). This suggests that algal growth rates were likely already near or at a maximum by this time, and coral physiologically impaired and susceptible to storm-induced damage, disease and algal overgrowth along the entire coast (Lapointe 1999; Bruno and others 2003; Wiedenmann and others 2013; Lapointe and others 2019) (Figure 1A, Supplement S.3). However, the persistent co-variation between eutrophication and *Diadema* grazing (Supplementary Figure S8) precluded identifying which of these two factors had a larger influence on coral reef composition and resilience at the time.

**Figure 6.**
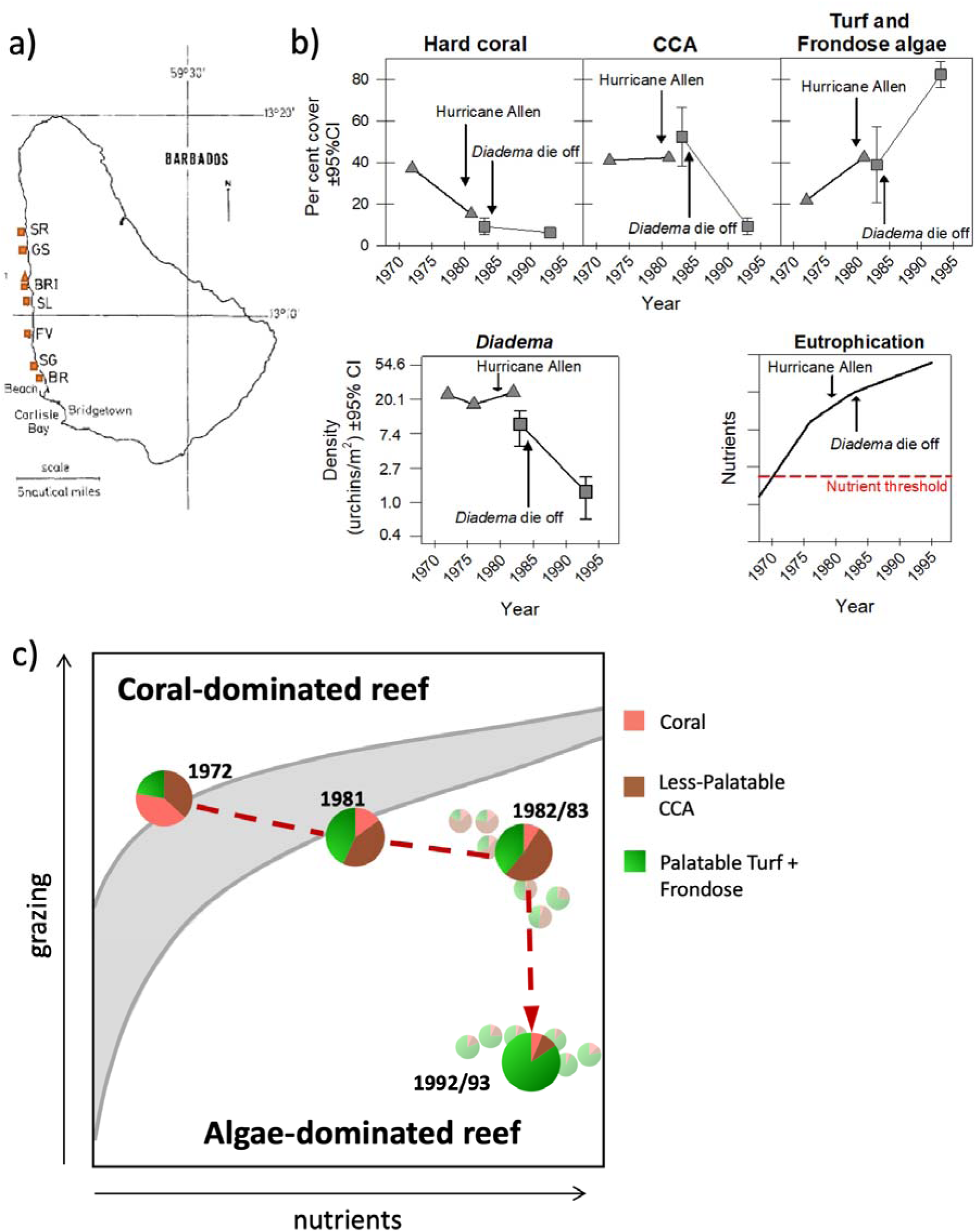
Changes in benthic composition of fringing reefs along the west coast of Barbados in space and time. A) Location of eight fringing reefs on the west coast of Barbados studied between 1972 and 1993. The triangle symbol shows the location of the North Bellairs reef, which was here used as baseline for 1972 (Stearn and others 1977) (pre-hurricane and pre-*Diadema* die off) and 1981 (Mah and Stearn 1986) (post-hurricane but pre-*Diadema* die off) whereas the square symbols represent seven other fringing reefs surveyed (Tomascik and Sander 1987a; Allard 1994), including South Bellairs reef (BRI), which is located only meters away from North Bellairs reef. From North to South: SR = Sandridge, GS = Greensleeves, North Bellairs Reef (triangle), BRI = Bellairs Research Institute (South Bellairs Reef), SL = Sandy Lane, FV = Fitts Village, SG = Spring Gardens, BR= Brighton; B) Temporal changes in the per cent cover of hard coral, CCA and turf + frondose algae in the fringing reefs, as well as changes in *Diadema* densities (95%CI when coastal average shown) and a schematic showing overall nutrient concentrations relative to eutrophication thresholds (see Supplement S.3 for individual nutrient measurements and thresholds); C) Mapping the trajectory of change of coral reefs along Barbados’ west coast within our theoretical multi-stressor framework. Approximate trajectory over time is represented by the dashed red line, with average reef composition at each time period displayed in pie charts (data described above and in (B)). In 1982/83 and 1992/93 surveys of multiple reefs were conducted along the coastline, following a gradient in eutrophication and grazing rates, with site-level variation shown by smaller transparent charts.

Following the 1983/84 *Diadema* die-off, Allard (1994) repeated Tomascik and Sander’s (1987a) surveys along the coastal eutrophication/grazing gradient (which persisted into the 1990’s, albeit with much lower grazing rates in an attempt to separate these effects (Hunte and Wittenberg 1992; Wittenberg and Hunte 1992; Allard 1994; Mann 1994)), and found that the overall loss of *Diadema* led to a subsequent increase in more-palatable turf and macroalgae along the entire west coast (Figure 6B). This was used to conclude that *Diadema* grazing was in fact the dominant driver of benthic composition along the west coast, ruling out the importance of the eutrophication gradient (Allard 1994). While the reduction in *Diadema* would have certainly altered benthic composition along the west coast of Barbados, Allard didn’t consider the already-severely-depleted state of coral by the early 1980s (Figure 6B) and we argue their conclusions instead reflected a shifting baseline rather than the full story of multi-stressor-induced change along the coast (Allard 1994; Jackson 2001b, 2001a). Indeed, the observed increase in palatable turf and frondose macroalgae appears to have been largely at the expense of less-palatable CCA rather than coral, reflecting this shifting baseline (Figure 6B). Additionally, by the 1990s the eutrophication/grazing gradient was weaker and no longer helped explain local variation in benthic composition between reefs as they were already in such a depleted state (Supplementary Figure S8). Grazing rates at this point were nearly homogeneous, the entire coastline was nutrient saturated, and the reefs were well-established as algae-dominated reefs. Collectively, these results highlight the context-dependency of our ability to detect multi-stressor effects on benthic composition in field studies and the importance of historical baselines.

Importantly, the patterns from Barbados’ west coast reefs support our theoretical predictions given changes to the multi-stressor landscape in both time and space, and do so at two spatial scales (i.e., along the entire coastline, and between local sites). Accordingly, we map these patterns in space and time as a trajectory of change within our multi-stressor framework (i.e., Figure 5), showing how the trajectory of benthic cover over time in Barbados can be explained as a function of both eutrophication and grazing (Figure 6C). Based on the timing of multi-stressor change described above, nutrients initially pushed the system through our “region of uncertainty” (likely also around the time of Hurricane Allen in 1980) and into an algae-dominated state (Figure 6C). The subsequent *Diadema* die-off was the final straw that further pushed the system well-into the algae-dominated region within our framework, and significantly altered the algae community composition (Figure 6C). While the specific placement and timing of this trajectory within our framework is not quantitative (grazing and nutrient thresholds are still debated and likely site-specific), we argue that this qualitative approach to viewing changes in multi-stressor impacts provides an important step towards a holistic understanding of coral reefs under global change, and allows us a way of piecing apart the roles of various drivers of change (e.g., the role of Barbados’ eutrophication/grazing gradient within this multi-stressor space). Here, we emphasize again the importance of baseline information to inform ongoing monitoring, and consideration of scale for disentangling the relative (and likely changing) influence of various drivers spatially and temporally.

## Discussion

Even with our relatively simple model used here, we have highlighted the important interactions between multi-stressor impacts, disturbance frequency, and life history strategies for the resilience of highly disturbed coral reef ecosystems. Immediately, our empirically-motivated coral reef model shows that changes in grazing, nutrients, and climate-driven coral mortality events (temperature stress, cyclonic storms) can independently invoke shifts to algae-dominated states and increase uncertainty in the face of global change. We also show that these three axes of global change operate synergistically to increase the likelihood of long transients and phase shifts to unwanted algal equilibrium states such that when all are conspiring together, we see even greater likelihood of deteriorated ecosystem structure and functioning. Our analysis found climate-driven mortality can drive shifts to algal domination even when in a moderately weakened state (i.e., nutrients and fishing are moderately high; Figure 2A ii) that would not be expected based on the deterministic skeleton. This novel theoretical result is related to recent theoretical literature on long transients (Hastings and others 2018, 2021), and importantly highlights the potential for unexpected results when considering noise in our mathematical models (Willcock and others 2023). Furthermore, our theoretical results are grounded in the geometry of our model and thus are highly general. That is, any combination of parameters that leads to the geometric configuration discussed throughout our manuscript (specifically, what we refer to as “moderate local impacts”) will lead to these results. Our results are intentionally general for this reason, but this of course implores us to use empirical approaches to find out the current state of reef resilience under system-specific parameterizations and multi-stressor impacts.

Our results also suggest the intriguing notion that the rise of different benthic r-strategists give us signatures of the dominant underlying stressor, matching a variety of empirical and conceptual accounts of benthic cover in coral reefs (Littler and Littler 1984; Lapointe 1997). Notably, fast-growing benthic r-strategists play important roles in highly disturbed coral reefs (Figure 3) and benthic composition is dependent on multi-stressors, particularly grazing-mediated algal selection that selects for differential life history (growth and palatability) strategies (Figures 1, 4)(Johnston and others 2022). This is consistent with experimental and observational studies (Sammarco 1980, 1982; Chiappone and others 2006), as well as theoretical concepts such as keystone predation theory (Leibold 1996). Here, less-palatable r2-strategists (e.g., CCA) garner a competitive advantage over more palatable fast-growth (r1) strategists when grazing rates are high, and vice versa. These life strategies are becoming even more important to consider in the face of global change and may offer us empirical insight into the mechanisms behind state shifts, including the potential for our “signatures of change” to act as early warning signals for coral collapse and decreasing resilience (related, see Nyström et al. (2008)). However, we note that further empirical research into underlying drivers is needed to establish more comprehensive signatures of multi-stressor impacts on benthic composition, as well as expand this model to include life history trade-offs within other taxa such as different macroalgae genera (Renken and Mumby 2009; Renken and others 2010; Bozec and others 2016; Briggs and others 2018), (hard and soft) corals (Zinke and others 2018; Toth and others 2019), sponges (González-Rivero and others 2011), and cyanobacteria (de Bakker and others 2017) for a more thorough representation of multi-stressor effects on benthic composition.

These signatures of multi-stressor impacts allow us to map changes in benthic composition within our predictive framework. Empirical evidence from Barbados shows promising consistency with our theoretical results and highlight the importance of accurate baseline data as well as monitoring multi-stressors simultaneously and with fine enough resolution to track changes and identify underlying drivers of change in coral reefs. Collectively, our empirical and theoretical results show that local stressors mediate coral reef resilience and can increase uncertainty in the face of climate-related mortality events, particularly as they are increasing in frequency and severity. Identifying underlying drivers of uncertainty are critical for management decisions, and highlight the importance of using precautionary approaches when success rates may be highly variable and unpredictable (Melbourne and Hastings 2009; White and others 2019).

Altogether, our generalizable approach harnessing the geometry of this model is an important pathway forward for developing a fundamental understanding of coral reef functioning under change as the mechanisms identified here ought to be common across many reefs and oceanic regions. However, it may not account for certain system-specific realities important for predicting the future state of individual reefs. For example, a precise parameterization would be necessary for assessing the relevant time scales of disturbances (and potential for recovery between them) critical for resilient reefs. Other factors such as spatial dynamics (Greiner and others 2022) and density-dependent mechanisms for corals or grazers (e.g., Allee effects) can have important implications for resilience against multi-stressors (disturbances, fishing pressure) and persistence, and may interact with each other (White and others 2021). Interactive effects with larger scale phenomena, as well as potential effects of other stressors, would be important to consider in future work.

While climate change is undoubtedly a massive threat to coral reefs, our results suggest that scientists and managers must simultaneously consider that local and regional management actions can facilitate at least some coral reef resilience (resistance to and recovery after disturbances). Here, we have shown how varying levels of multi-stressors can lead to very different trajectories of change in coral reefs (Figure 6), and would in turn have drastically different responses to singular management actions (e.g., Figure 2B,C). In times of rapid and drastic global change, research and management need to move away from isolating individual impacts and develop ways to address multiple stressors where nonlinear interactions are likely the rule, not the exception. This way we can manage for ecosystem resilience and sustainability in the face of uncertainty (Anthony and others 2015; Roberts and others 2017; Mcleod and others 2019). We also need effective management so that peoples’ livelihoods are not unnecessarily limited or sacrificed under the guise of conservation (e.g., through fishing bans or expanded no-take zones, when in reality there may be a much more nuanced problem that must be acknowledged (Aronson and Precht 2006). Ecosystem stability and sustainability ultimately means protecting important ecosystem services necessary for food and livelihood security and overall human wellbeing.

## Acknowledgements

This project was funded through a Canada First Research Excellence Fund project “Food from Thought” to KSM and CB was supported by a NSERC CGS-D.

We would like to thank Dr. Gabriel Gellner for his helpful discussions early on when designing this study and regarding the theoretical results and Dr. Jacob Allgeier for useful feedback on a previous draft.

## Author Contributions

All authors contributed to the development of ideas through workshops and discussions. CB and KSM designed the theoretical model and CB created the figures. Data collection from Barbados and creation of Figure 6 was done by HV. CB and KSM wrote the initial draft of the manuscript and all authors contributed to editing subsequent revisions.

## Notes

https://github.com/carlingbieg/corals

### Competing Interest Statement

The authors have declared no competing interest.

### Summary of Updates

Modest revisions as a result of peer review, including one figure change from original (Figures 6 and 7 combined).

## Literature Cited

Abelson A. 2020. Are we sacrificing the future of coral reefs on the altar of the “climate change” narrative? Browman H, editor. ICES J Mar Sci 77:40–5. https://academic.oup.com/icesjms/article/77/1/40/5673597

Allard P. 1994. Changes in Coral Community Structure in Barbados: Effects of Eutrophication and Reduced Grazing Pressure.

Andréfouët S, Mumby P, McField M, Hu C, Muller-Karger F. 2002. Revisiting coral reef connectivity. Coral Reefs 21:43–8. 10.1007/s00338-001-0199-0

Anthony KRN, Marshall PA, Abdulla A, Beeden R, Bergh C, Black R, Eakin CM, Game ET, Gooch M, Graham NAJ, Green A, Heron SF, van Hooidonk R, Knowland C, Mangubhai S, Marshall N, Maynard JA, Mcginnity P, Mcleod E, Mumby PJ, Nyström M, Obura D, Oliver J, Possingham HP, Pressey RL, Rowlands GP, Tamelander J, Wachenfeld D, Wear S. 2015. Operationalizing resilience for adaptive coral reef management under global environmental change. Glob Chang Biol 21:48–61.

Anthony KRN, Maynard JA, Diaz-Pulido G, Mumby PJ, Marshall PA, Cao L, Hoegh-Guldberg O. 2011. Ocean acidification and warming will lower coral reef resilience. Glob Chang Biol 17:1798–808.

Arias-González JE, Fung T, Seymour RM, Garza-Pérez JR, Acosta-González G, Bozec Y-M, Johnson CR. 2017. A coral-algal phase shift in Mesoamerica not driven by changes in herbivorous fish abundance. Bernardi G, editor. PLoS One 12:e0174855. https://dx.plos.org/10.1371/journal.pone.0174855. Last accessed 11/11/2020

Aronson RB, Precht WF. 2001. White-band disease and the changing face of Caribbean coral reefs. Hydrobiologia 460:25–38.

Aronson RB, Precht WF. 2006. Conservation, precaution, and Caribbean reefs. Coral Reefs 25:441–50. http://link.springer.com/10.1007/s00338-006-0122-9. Last accessed 11/11/2020

de Bakker DM, van Duyl FC, Bak RPM, Nugues MM, Nieuwland G, Meesters EH. 2017. 40 Years of benthic community change on the Caribbean reefs of Curaçao and Bonaire: the rise of slimy cyanobacterial mats. Coral Reefs 36:355–67.

Ban SS, Graham NAJ, Connolly SR. 2014. Evidence for multiple stressor interactions and effects on coral reefs. Glob Chang Biol 20:681–97.

Bell PRF, Elmetri I, Lapointe BE. 2014. Evidence of Large-Scale Chronic Eutrophication in the Great Barrier Reef: Quantification of Chlorophyll a Thresholds for Sustaining Coral Reef Communities. Ambio 43:361–76. http://link.springer.com/10.1007/s13280-013-0443-1

Bell PRF, Tomascik T. 1994. The demise of the fringing coral reefs of Barbados and of regions in the Great Barrier Reef (GBR) lagoon - impacts of eutrophication. Proc Colloq Glob Asp coral reefs, Miami, 1993:319–25.

Blackwood JC, Hastings A, Mumby PJ. 2011. A model-based approach to determine the long-term effects of multiple interacting stressors on coral reefs. Ecol Appl 21:2722–33. http://doi.wiley.com/10.1890/10-2195.1

Blackwood JC, Hastings A, Mumby PJ. 2012. The effect of fishing on hysteresis in Caribbean coral reefs. Theor Ecol 5:105–14.

Blackwood JC, Okasaki C, Archer A, Matt EW, Sherman E, Montovan K. 2018. Modeling alternative stable states in Caribbean coral reefs. Nat Resour Model 31:1–17.

Bozec YM, O’Farrell S, Bruggemann JH, Luckhurst BE, Mumby PJ. 2016. Tradeoffs between fisheries harvest and the resilience of coral reefs. Proc Natl Acad Sci U S A 113:4536–41.

Brandl SJ, Rasher DB, Côté IM, Casey JM, Darling ES, Lefcheck JS, Duffy JE. 2019. Coral reef ecosystem functioning: eight core processes and the role of biodiversity. Front Ecol Environ 17:445–54.

Briggs CJ, Adam TC, Holbrook SJ, Schmitt RJ. 2018. Macroalgae size refuge from herbivory promotes alternative stable states on coral reefs. PLoS One 13:1–14.

Bruno JF, Côté IM, Toth LT. 2019. Climate Change, Coral Loss, and the Curious Case of the Parrotfish Paradigm: Why Don’t Marine Protected Areas Improve Reef Resilience? Ann Rev Mar Sci 11:307–34.

Bruno JF, Petes LE, Harvell CD, Hettinger A. 2003. Nutrient enrichment can increase the severity of coral diseases. Ecol Lett 6:1056–61.

Burkepile DE, Hay ME. 2006. Herbivore vs. nutrient control of marine primary producers: Context-dependent effects. Ecology 87:3128–39.

Ceccarelli DM, Jones GP, McCook LJ. 2011. Interactions between herbivorous fish guilds and their influence on algal succession on a coastal coral reef. J Exp Mar Bio Ecol 399:60–7. http://www.sciencedirect.com/science/article/pii/S002209811100044X

Chiappone M, Swanson D, Miller S, Science M. 2006. One-year Response of Florida Keys Patch Reef Communities to Translocation of Long-spined Sea Urchins (Diadema antillarum). Sites J 20Th Century Contemp French Stud:319–44.

Cline TJ, Allgeier JE. 2022. Fish community structure and dynamics are insufficient to mediate coral resilience. Nat Ecol Evol 6:1700–9.

Cote IM, Darling ES, Brown CJ. 2016. Interactions among ecosystem stressors and their importance in conservation. Proc R Soc B Biol Sci 283:1–9.

DeCarlo TM, Gajdzik L, Ellis J, Coker DJ, Roberts MB, Hammerman NM, Pandolfi JM, Monroe AA, Berumen ML. 2020. Nutrient-supplying ocean currents modulate coral bleaching susceptibility. Sci Adv 6:1–8.

Done TJ. 1992. Phase shifts in coral reef communities and their ecological significance. Hydrobiologia 247:121–32.

Donovan MK, Adam TC, Shantz AA, Speare KE, Munsterman KS, Rice MM, Schmitt RJ, Holbrook SJ, Burkepile DE. 2020. Nitrogen pollution interacts with heat stress to increase coral bleaching across the seascape. Proc Natl Acad Sci U S A 117:5351–7.

Donovan MK, Burkepile DE, Kratochwill C, Shlesinger T, Sully S, Oliver TA, Hodgson G, Freiwald J, van Woesik R. 2021. Local conditions magnify coral loss after marine heatwaves. Science (80-) 372:977–80.

Fung T, Seymour RM, Johnson CR. 2011. Alternative stable states and phase shifts in coral reefs under anthropogenic stress. Ecology 92:967–82. http://doi.wiley.com/10.1890/10-0378.1

Gardner TA, Côté IM, Gill JA, Grant A, Watkinson AR. 2003. Long-term region-wide declines in Caribbean corals. Science (80-) 301:958–60.

Gill DA, Oxenford HA, Turner RA, Schuhmann PW. 2019. Making the most of data-poor fisheries: Low cost mapping of small island fisheries to inform policy. Mar Policy 101:198–207.

González-Rivero M, Yakob L, Mumby PJ. 2011. The role of sponge competition on coral reef alternative steady states. Ecol Modell 222:1847–53. https://linkinghub.elsevier.com/retrieve/pii/S0304380011001438

Greiner A, S. Darling E, Fortin MJ, Krkošek M. 2022. The combined effects of dispersal and herbivores on stable states in coral reefs. Theor Ecol 15:321–35. 10.1007/s12080-022-00546-w

Guan Y, Hohn S, Wild C, Merico A. 2020. Vulnerability of global coral reef habitat suitability to ocean warming, acidification and eutrophication. Glob Chang Biol 26:5646–60.

Gurney GG, Melbourne-Thomas J, Geronimo RC, Aliño PM, Johnson CR. 2013. Modelling coral reef futures to inform management: Can reducing local-scale stressors conserve reefs under climate change? PLoS One 8:1–17.

Halpern BS, Frazier M, Afflerbach J, Lowndes JS, Micheli F, O’Hara C, Scarborough C, Selkoe KA. 2019. Recent pace of change in human impact on the world’s ocean. Sci Rep 9:1–8.

Harborne AR, Rogers A, Bozec Y-M, Mumby PJ. 2017. Multiple Stressors and the Functioning of Coral Reefs. Ann Rev Mar Sci 9:445–68.

Hastings A, Abbott KC, Cuddington K, Francis T, Gellner G, Lai YC, Morozov A, Petrovskii S, Scranton K, Zeeman M Lou. 2018. Transient phenomena in ecology. Science (80-) 361.

Hastings A, Abbott KC, Cuddington K, Francis TB, Lai YC, Morozov A, Petrovskii S, Zeeman M Lou. 2021. Effects of stochasticity on the length and behaviour of ecological transients. J R Soc Interface 18.

Higgins K, Hastings A, Sarvela JN, Botsford LW. 1997. Stochastic Dynamics and Deterministic Skeletons: Population Behavior of Dungeness Crab. Science (80-) 276:1431–5. http://science.sciencemag.org/. Last accessed 11/11/2020

Hixon MA, Brostoff WN. 1996. Succession and herbivory: Effects of differential fish grazing on hawaiian coral-reef algae. Ecol Monogr 66:67–90. https://esajournals.onlinelibrary.wiley.com/doi/full/10.2307/2963481. Last accessed 11/11/2020

Houk P, Castro F, McInnis A, Rucinski M, Starsinic C, Concepcion T, Manglona S, Salas E. 2022. Nutrient thresholds to protect water quality, coral reefs, and nearshore fisheries. Mar Pollut Bull 184:114144.

Hughes TP. 1994. Catastrophes, Phase Shifts, and Large-Scale Degradation of a Caribbean Coral Reef. Science (80-) 265:1547–51. https://www.sciencemag.org/lookup/doi/10.1126/science.265.5178.1547

Hughes TP, Barnes ML, Bellwood DR, Cinner JE, Cumming GS, Jackson JBC, Kleypas J, Van De Leemput IA, Lough JM, Morrison TH, Palumbi SR, Van Nes EH, Scheffer M. 2017. Coral reefs in the Anthropocene. Nature 546:82–90.

Hughes TP, Connell JH. 1999. Multiple stressors on coral reefs: A long-term perspective. Limnol Oceanogr 44:932–40.

Hughes TP, Rodrigues MJ, Bellwood DR, Ceccarelli D, Hoegh-Guldberg O, McCook L, Moltschaniwskyj N, Pratchett MS, Steneck RS, Willis B. 2007. Phase Shifts, Herbivory, and the Resilience of Coral Reefs to Climate Change. Curr Biol 17:360–5.

Hunte W, Wittenberg M. 1992. Effects of eutrophication and sedimentation on juvenile corals II. Settlement. Mar Biol 114:625–31.

Jackson J, Donovan M, Cramer K, Lam V. 2014. Status and trends of Caribbean coral reefs: 1970-2012. Gland, Switzerland http://pubs.er.usgs.gov/publication/70115405

Jackson JBC. 2001a. Historical Overfishing and the Recent Collapse of Coastal Ecosystems. Science (80-) 293:629–37. www.sciencemag.org. Last accessed 01/03/2021

Jackson JBC. 2001b. What was natural in the coastal oceans? Proc Natl Acad Sci 98:5411–8. www.pnas.orgcgidoi10.1073pnas.091092898. Last accessed 01/03/2021

Johnston EL, Clark GF, Bruno JF. 2022. The speeding up of marine ecosystems. Clim Chang Ecol 3:100055.

Jouffray JB, Nyström M, Norström A V., Williams ID, Wedding LM, Kittinger JN, Williams GJ. 2015. Identifying multiple coral reef regimes and their drivers across the hawaiian archipelago. Philos Trans R Soc B Biol Sci 370:1–8.

Lapointe BE. 1997. Nutrient thresholds for bottom-up control of macroalgal blooms on coral reefs in Jamaica and southeast Florida. Limnol Oceanogr 42:1119–31.

Lapointe BE. 1999. Simultaneous top-down and bottom-up forces control macroalgal blooms on coral reefs (Reply to the comment by Hughes et al.). Limnol Oceanogr 44:1586–92. http://doi.wiley.com/10.4319/lo.1999.44.6.1586

Lapointe BE, Brewton RA, Herren LW, Porter JW, Hu C. 2019. Nitrogen enrichment, altered stoichiometry, and coral reef decline at Looe Key, Florida Keys, USA: a 3-decade study. Mar Biol 166:1–31. 10.1007/s00227-019-3538-9

Lapointe BE, Burkholder JM, Van Alstyne KL. 2018. Harmful Macroalgal Blooms in a Changing World: Causes, Impacts, and Management. In: Harmful Algal Blooms. Chichester, UK: John Wiley & Sons, Ltd. pp 515–60. http://doi.wiley.com/10.1002/9781118994672.ch15

Leibold MA. 1996. A Graphical Model of Keystone Predators in Food Webs: Trophic Regulation of Abundance, Incidence, and Diversity Patterns in Communities. Am Nat 147:784–812.

Lessios HA. 1988. Mass mortality of Diadema antillarum in the Caribbean: what have we learned? Annu Rev Ecol Syst Vol 19:371–93.

Lewis, John B. 1960. The coral reefs and coral communities of Barbados, W.I. Can J Zool 38:1133–52.

Littler M, Littler D. 1984.Models of tropical reef biogenesis: the contribution of algae.

MacNeil MA, Mellin C, Matthews S, Wolff NH, McClanahan TR, Devlin M, Drovandi C, Mengersen K, Graham NAJ. 2019. Water quality mediates resilience on the Great Barrier Reef. Nat Ecol Evol 3:620–7. 10.1038/s41559-019-0832-3

Mah AJ, Stearn CW. 1986. The effect of Hurricane Allen on the Bellairs fringing reef, Barbados. Coral Reefs 4:169–76. 10.1007/BF00427938

Mann GS. 1994.Distribution, abundance and life history of the reef coral Favia fragrum (Esper) in Barbados: effects of eutrophication and of the black sea urchin Diadema antillarum (Philippi).

Mcclanahan TR. 1997.Primary succession of coral-reef algae: Differing patterns on fished versus unfished reefs.

McClanahan TR, Sala E, Stickels PA, Cokos BA, Baker AC, Starger CJ, Jones IV SH. 2003. Interaction between nutrients and herbivory in controlling algal communities and coral condition on Glover’s Reef, Belize. Mar Ecol Prog Ser 261:135–47.

Mcleod E, Anthony KRN, Mumby PJ, Maynard J, Beeden R, Graham NAJ, Heron SF, Hoegh-Guldberg O, Jupiter S, MacGowan P, Mangubhai S, Marshall N, Marshall PA, McClanahan TR, Mcleod K, Nyström M, Obura D, Parker B, Possingham HP, Salm R V., Tamelander J. 2019. The future of resilience-based management in coral reef ecosystems. J Environ Manage 233:291–301. 10.1016/j.jenvman.2018.11.034

McManus JW, Meñez LAB, Kesner-Reyes KN, Vergara SG, Ablan MC. 2000. Coral reef fishing and coral-algal phase shifts: implications for global reef status. ICES J Mar Sci 57:572–8. https://academic.oup.com/icesjms/article-lookup/doi/10.1006/jmsc.2000.0720

Melbourne BA, Hastings A. 2009. Highly Variable Spread Rates in Replicated Biological Invasions: Fundamental Limits to Predictability. Science (80-) 325:1536–9. https://www.science.org/doi/10.1126/science.1176138

Meyer K, Hoyer-Leitzel A, Iams S, Klasky I, Lee V, Ligtenberg S, Bussmann E, Zeeman M Lou. 2018. Quantifying resilience to recurrent ecosystem disturbances using flow–kick dynamics. Nat Sustain 1:671–8. 10.1038/s41893-018-0168-z

Morozov A, Abbott K, Cuddington K, Francis T, Gellner G, Hastings A, Lai YC, Petrovskii S, Scranton K, Zeeman M Lou. 2020. Long transients in ecology: Theory and applications. Phys Life Rev 1. 10.1016/j.plrev.2019.09.004

Mumby PJ. 2006. The impact of exploiting grazers (Scaridae) on the dynamics of Caribbean coral reefs. Ecol Appl 16:747–69.

Mumby PJ. 2017. Embracing a world of subtlety and nuance on coral reefs. Coral Reefs 36:1003–11.

Mumby PJ, Hastings A, Edwards HJ. 2007. Thresholds and the resilience of Caribbean coral reefs. Nature 450:98–101.

Mumby PJ, Hedley JD, Zychaluk K, Harborne AR, Blackwell PG. 2006. Revisiting the catastrophic die-off of the urchin Diadema antillarum on Caribbean coral reefs: Fresh insights on resilience from a simulation model. Ecol Modell 196:131–48. https://linkinghub.elsevier.com/retrieve/pii/S0304380005006174

Mumby PJ, Steneck RS. 2008. Coral reef management and conservation in light of rapidly evolving ecological paradigms. Trends Ecol Evol 23:555–63. http://www.sciencedirect.com/science/article/pii/S0169534708002504

Mumby PJ, Wolff NH, Bozec YM, Chollett I, Halloran P. 2014. Operationalizing the resilience of coral reefs in an era of climate change. Conserv Lett 7:176–87.

Muthukrishnan R, Fong P. 2014. Multiple anthropogenic stressors exert complex, interactive effects on a coral reef community. Coral Reefs 33:911–21.

Norström A V., Nyström M, Jouffray JB, Folke C, Graham NAJ, Moberg F, Olsson P, Williams GJ. 2016. Guiding coral reef futures in the Anthropocene. Front Ecol Environ 14:490–8.

Norström A V., Nyström M, Lokrantz J, Folke C. 2009. Alternative states on coral reefs: Beyond coral-macroalgal phase shifts. Mar Ecol Prog Ser 376:293–306.

Nyström M, Graham NAJ, Lokrantz J, Norström A V. 2008. Capturing the cornerstones of coral reef resilience: Linking theory to practice. Coral Reefs 27:795–809.

Pendleton LH, Hoegh-Guldberg O, Langdon C, Comte A. 2016. Multiple stressors and ecological complexity require a new approach to coral reef research. Front Mar Sci 3:1–5.

Renken H, Mumby PJ. 2009. Modelling the dynamics of coral reef macroalgae using a Bayesian belief network approach. Ecol Modell 220:1305–14. https://linkinghub.elsevier.com/retrieve/pii/S0304380009001525

Renken H, Mumby PJ, Matsikis I, Edwards HJ. 2010. Effects of physical environmental conditions on the patch dynamics of Dictyota pulchella and Lobophora variegata on Caribbean coral reefs. Mar Ecol Prog Ser 403:63–74.

Roberts M, Hanley N, Williams S, Cresswell W. 2017. Terrestrial degradation impacts on coral reef health: Evidence from the Caribbean. Ocean Coast Manag 149:52–68. 10.1016/j.ocecoaman.2017.09.005

Robinson JPW, Wilson SK, Graham NAJ. 2019. Abiotic and biotic controls on coral recovery 16 years after mass bleaching. Coral Reefs 38:1255–65. 10.1007/s00338-019-01831-7

Sammarco PW. 1980. Diadema and its relationship to coral spat mortality: grazing, competition, and biological disturbance. J Exp Mar Bio Ecol 45:245–72.

Sammarco PW. 1982. Effects of grazing by Diadema antillarum Philippi (Echinodermata: Echinoidea) on algal diversity and community structure. J Exp Mar Bio Ecol 65:83–105.

Sander F, Moore E. 1979. Significance of ammonia in determining the N:P ratio of the sea water off Barbados, West Indies. Mar Biol 55:17–21.

Scheffer M, Barrett S, Carpenter SR, Folke C, Green AJ, Holmgren M, Hughes TP, Kosten S, Van De Leemput IA, Nepstad DC, Van Nes EH, Peeters ETHM, Walker B. 2015. Creating a safe operating space for iconic ecosystems: Manage local stressors to promote resilience to global change. Science (80-) 347:1317–9.

Shantz AA, Burkepile DE. 2014. Context-dependent effects of nutrient loading on the coral-algal mutualism. Ecology 95:1995–2005.

Smith J, Smith C, Hunter C. 2001. An experimental analysis of the effects of herbivory and nutrient enrichment on benthic community dynamics on a Hawaiian reef. Coral Reefs 19:332–42. 10.1007/s003380000124

Stearn CW, Scoffin TP, Martindale W. 1977. Calcium carbonate budget of a fringing reef on the west coast of Barbados Part I- zonation and productivity. Bull Mar Sci 27:479–510.

Steffen W, Broadgate W, Deutsch L, Gaffney O, Ludwig C. 2015. The trajectory of the anthropocene: The great acceleration. Anthr Rev 2:81–98.

Steneck RS, Arnold SN, Mumby PJ. 2014. Experiment mimics fishing on parrotfish: Insights on coral reef recovery and alternative attractors. Mar Ecol Prog Ser 506:115–27.

Suchley A, Alvarez-Filip L. 2018. Local human activities limit marine protection efficacy on Caribbean coral reefs. Conserv Lett 11:e12571. http://doi.wiley.com/10.1111/conl.12571. Last accessed 11/11/2020

Thampi VA, Anand M, Bauch CT. 2018. Socio-ecological dynamics of Caribbean coral reef ecosystems and conservation opinion propagation. Sci Rep 8:1–11. 10.1038/s41598-018-20341-0

Tilman D, May RM, Lehman CL, Nowak MA. 1994. Habitat destruction and the extinction debt. Nature 371:65–6. http://www.nature.com/articles/371065a0

Tomascik T. 1990. Growth rates of two morphotypes of Montastrea annularis along a eutrophication gradient, Barbados, W.I. Mar Pollut Bull 21:376–81.

Tomascik T. 1991. Settlement patterns od Caribbean scleractinian corals on artificial substrata along a eutrophication gradient, Barbados, West Indies. Mar Ecol Prog Ser 77:261–9.

Tomascik T, Sander F. 1985. Effects of eutrophication on reef-building corals I. Growth rate of the reef-building coral Montastrea annularis. Mar Biol 87.

Tomascik T, Sander F. 1987a. Effects of eutrophication on reef-building corals - II. Structure of scleractinian coral communities on fringing reefs, Barbados, West Indies. Mar Biol 94:53–75.

Tomascik T, Sander F. 1987b. Effects of eutrophication on reef-building corals - III. Reproduction of the reef-building coral Porites porites. Mar Biol 94:77–94.

Tosic M, Bonnell RB, Dutilleul P, Oxenford HA. 2008. Runoff Water Quality, Landuse and Environmental Impacts on the Bellairs Fringing Reef, Barbados. In: Remote Sensing and Geospatial Technologies for Coastal Ecosystem Assessment and Management. Springer Berlin Heidelberg. pp 521–53. https://link.springer.com/chapter/10.1007/978-3-540-88183-4_22. Last accessed 26/02/2021

Toth LT, Stathakopoulos A, Kuffner IB, Ruzicka RR, Colella MA, Shinn EA. 2019. The unprecedented loss of Florida’s reef-building corals and the emergence of a novel coral-reef assemblage. Ecology 100:1–14.

Vega Thurber RL, Burkepile DE, Fuchs C, Shantz AA, Mcminds R, Zaneveld JR. 2014. Chronic nutrient enrichment increases prevalence and severity of coral disease and bleaching. Glob Chang Biol 20:544–54.

Wang L, Shantz AA, Payet JP, Sharpton TJ, Foster A, Burkepile DE, Thurber RV. 2018. Corals and their microbiomes are differentially affected by exposure to elevated nutrients and a natural thermal anomaly. Front Mar Sci 5:1–16.

Wear SL. 2016. Missing the boat: Critical threats to coral reefs are neglected at global scale. Mar Policy 74:153–7. 10.1016/j.marpol.2016.09.009

Wear SL, Thurber RV. 2015. Sewage pollution: Mitigation is key for coral reef stewardship. Ann N Y Acad Sci 1355:15–30.

White ER, Baskett ML, Hastings A. 2021. Catastrophes, connectivity and Allee effects in the design of marine reserve networks. Oikos 130:366–76.

White ER, Cox K, Melbourne BA, Hastings A. 2019. Success and failure of ecological management is highly variable in an experimental test. Proc Natl Acad Sci U S A 116:23169–73.

Wiedenmann J, D’Angelo C, Smith EG, Hunt AN, Legiret FE, Postle AD, Achterberg EP. 2013. Nutrient enrichment can increase the susceptibility of reef corals to bleaching. Nat Clim Chang 3:160–4.

Willcock S, Cooper GS, Addy J, Dearing JA. 2023. Earlier collapse of Anthropocene ecosystems driven by multiple faster and noisier drivers. Nat Sustain.

Wittenberg M, Hunte W. 1992. Effects of eutrophication and sedimentation on juvenile corals I. Abundance, mortality and community structure. Mar Biol 112:131–8.

Wooldridge SA. 2009. Water quality and coral bleaching thresholds: Formalising the linkage for the inshore reefs of the Great Barrier Reef, Australia. Mar Pollut Bull 58:745–51. 10.1016/j.marpolbul.2008.12.013

Wooldridge SA, Done TJ. 2012. water quality can ameliorate effects Improved of climate on corals change. 19:1492–9.

Zaneveld JR, Burkepile DE, Shantz AA, Pritchard CE, McMinds R, Payet JP, Welsh R, Correa AMS, Lemoine NP, Rosales S, Fuchs C, Maynard JA, Thurber RV. 2016. Overfishing and nutrient pollution interact with temperature to disrupt coral reefs down to microbial scales. Nat Commun 7:1–12.

Zinke J, Gilmour JP, Fisher R, Puotinen M, Maina J, Darling E, Stat Michael, Richards ZT, Mcclanahan TR, Beger M, Moore Cordelia, Graham NAJ, Feng M, Hobbs J-PA, Evans SN, Field Stuart, Shedrawi G, Russ, Babcock C, Wilson SK, Macarthur CT. 2018. Gradients of disturbance and environmental conditions shape coral community structure for south-eastern Indian Ocean reefs. Divers Distrib:24.

