## Supplement for "Towards a multi-stressor theory for coral reefs in a changing world"

**Supplementary Material for manuscript: “Towards a multi-stressor theory for coral reefs in a changing world”**

**S.1 Model geometry and theoretical results**

***Geometry of model – phase space and the effect of parameters on isoclines and equilibria***

Recall our coral reef model, a modified version of that developed by Mumby et al. (2007):

  $\frac{dM}{dt}=M(aC\left( \frac{N_{t}}{\left( N_{0}+N_{t} \right)} \right)- \frac{g}{\left( M+B \right)} + yB)$	(1)

$\frac{dC}{dt}=C(r\left( 1-cN_{t} \right)B-m-aM\left( \frac{N_{t}}{\left( N_{0}+N_{t} \right)} \right))$	(2)

where M represents macroalgae, C represents coral, and B represents other benthic r-strategists that instantaneously fill empty space left by dead coral. Since B is assumed to instantaneously fill any space (r-strategists act on much faster time scales relative to the other state variables), so B = 1 – M – C and the model can therefore be reduced to two dimensions.

To solve for the Macroalgae isocline (shown as solid line in Supplementary Figure C1 below), we set dM/dt = 0 (Equation 1) and solve for M. Therefore,

$\frac{dM}{dt}=0: M=1-C + \frac{aC}{y}\left( \frac{N_{t}}{\left( N_{0}+N_{t} \right)} \right)- \frac{g}{y(1-C)}$	(3)

Similarly, we can solve for the Coral isocline (shown as dashed line in Supplementary Figure C1) such that dC/dt = 0 (Equation 2) and solve for M. In this case,

$\frac{dC}{dt}=0: M=\frac{\left( N_{0}+N_{t} \right)\left( m-r(1-cN_{t} \right)(1-C))}{r\left( N_{0}+N_{t} \right)\left( cN_{t}-1 \right)-aN_{t}}$	(4)

These isoclines (3 and 4) are displayed in Supplementary Figure C1 and show the equilibria structure (stable and unstable equilibria) when in a bistable configuration. Note that these equilibria of course change as various parameters (notably including both *g* and *N_t_*) can drive bifurcations.

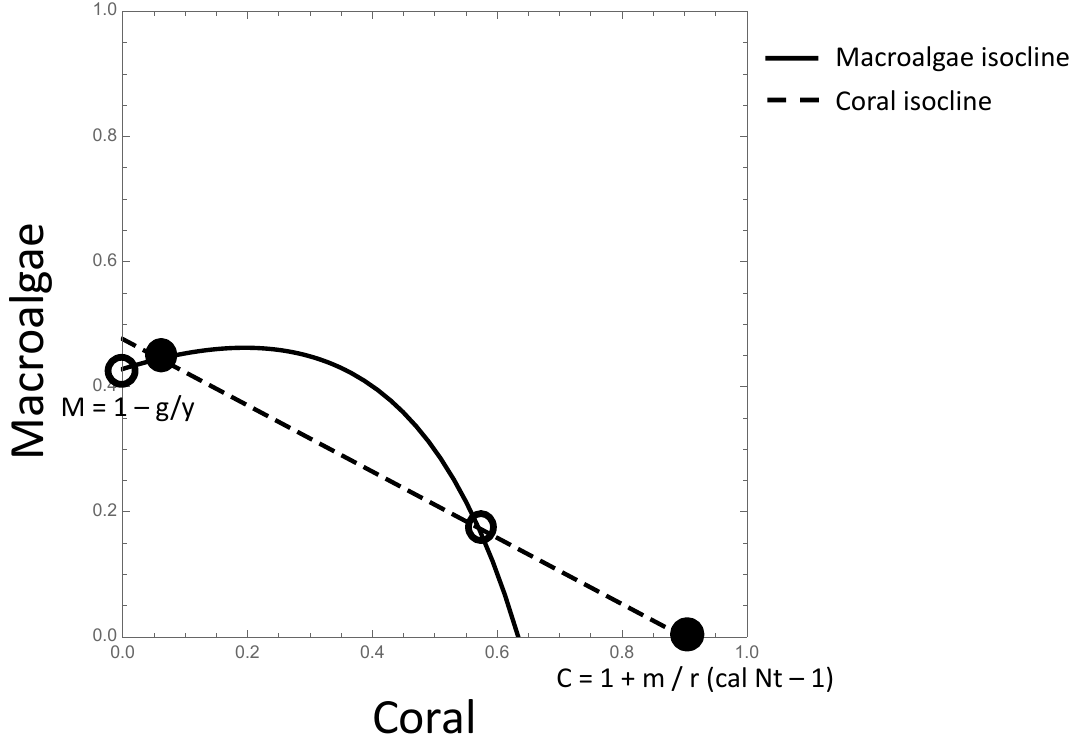
**Supplementary Figure S1.** Isocline structure of our nutrient-dependent extension of Mumby et al.(Mumby et al. 2007), showing the macroalgae isocline (solid line), coral isocline (dashed line), stable equilibria (full circles) and unstable equilibria (open circles) in a bistable configuration.

We can immediately tell by the isocline equations (3 and 4) that both grazing (*g*) and nutrients (*N_t_*) alter the geometry of the model. Specifically, they both act in qualitatively similar ways to shift the isoclines (*g* and *N_t_* are both in the numerator for the Macroalgae isocline (3) but note differing signs, +/-) and drive a saddle node bifurcation. Along with other parameters, nutrients additionally change the slope and intersects of the Coral isocline (4) with the axes, and change where the isoclines intersect (i.e., the equilibria). Clearly, if both stressors are changed simultaneously, these effects (including bifurcations) will be seen even more quickly. Supplementary Figure C2 shows the separate effects of grazing and nutrients on the isocline structure.

While the interior equilibria are too complex to show symbolically, we can importantly see how grazing and nutrients influence the axial solutions shown in Supplementary Figure C1 above. Note that the Coral axial solution is a stable equilibrium, and defines the benthic composition when the system is at a coral-dominated state (M = 0; B = 1-C), and importantly depends on nutrient loading.

Initially, the bistable configuration with one stable axial solution and one stable interior solution (as shown in Supplementary Figure C1) means that we can have a coral-dominated state (M=0) and an algae-dominated state where coral is low but not 0. However, the stable interior and unstable M-axial (C=0) solutions quickly undergo a transcritical bifurcation with increasing stressors (nutrients and/or grazing), meaning that the Macroalgae axial solution becomes stable at even moderate levels of human impact. In this case, the system remains bistable, however there is no possibility for coexistence of M and C. This axial solution importantly determines the equilibrium value for a Macroalgae dominated state (C = 0), and is determined by the level of grazing suppression imposed on Macroalgae. Note that in both cases the remaining benthic cover is composed of other benthic r-strategists, such as turf algae and CCA (B = 1 – M – C).

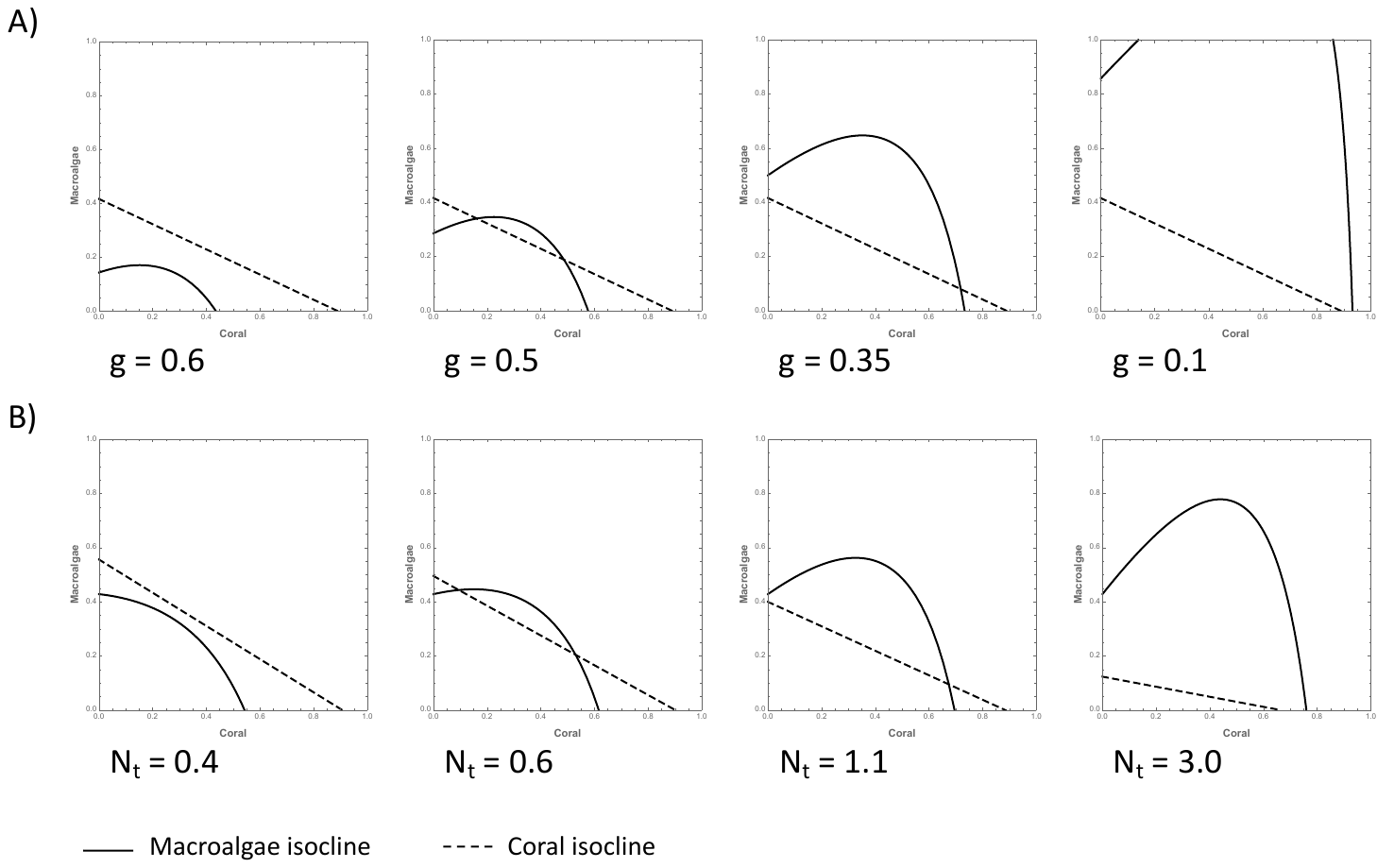
**Supplementary Figure S2.** Isocline structure and the effect of (A) decreasing grazing, and (B) increasing nutrients while holding all other parameters constant. *a* = 2.3, *y* = 0.7, *r* = 1.8, *m* = 0.15, *N*_0_ = 0.5, *c* = 0.25, *g* (when not varied) = 0.4 and *N*_t_ (when not varied) = 1.0.

**S.2 Transient dynamics and the effect of noise**

***Long transients and extinction debt in the deterministic model***

Phaseplane geometry and initial values in certain configurations can lead to long transients, even in deterministic simulations. Supplementary Figure C3 shows one example of a long transient. In this case coral cover initially increases, approaching the unstable saddle point, before crossing the M isocline and being pulled towards the axial solution on the M axis (C=0). Without looking at the basins of attraction to each stable equilibria, one might think that coral would recover based on its initial trajectory (after a single pulse perturbation), however in this case it does not reach this equilibrium. This is an example of potential extinction debt in bistable configurations, such that coral is doomed to eventual extinction without any additional mortality or stressors, but this outcome may be masked for some time.

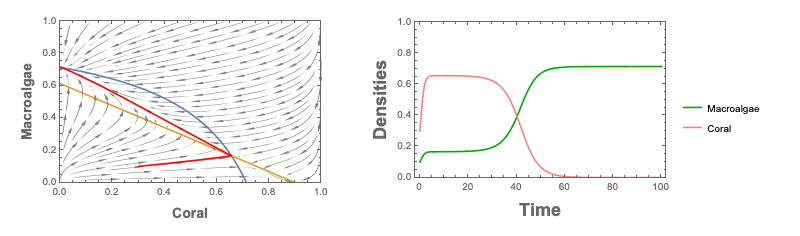
**Supplementary Figure S3.** Relatively long transients in a deterministic simulation. A) isocline geometry (dM/dt=0 in blue and dC/dt=0 in yellow) and trajectory through phase space (red line). B) the same trajectory displayed in (a) but shown as a time series, displaying how coral initially increases and macroalgae is suppressed for some time, before coral eventually goes extinct. Here, *y* = 0.7, *r* = 1.8, *m* = 0.15, *N*_0_ = 0.5, *c* = 0.25, g=0.2, a=1.2, Nt=0.69.

***Incorporating noise through discrete perturbations (climate-driven mortality events)***

Incorporating noise through repeated pulse perturbations to coral cover (-50%), can amplify these transient dynamics such that we see extremely long transients under certain conditions, particularly under moderate human impact scenarios (where the system gets entangled near a ghost saddle). Supplementary Figure C4 shows an example of this long transient scenario.

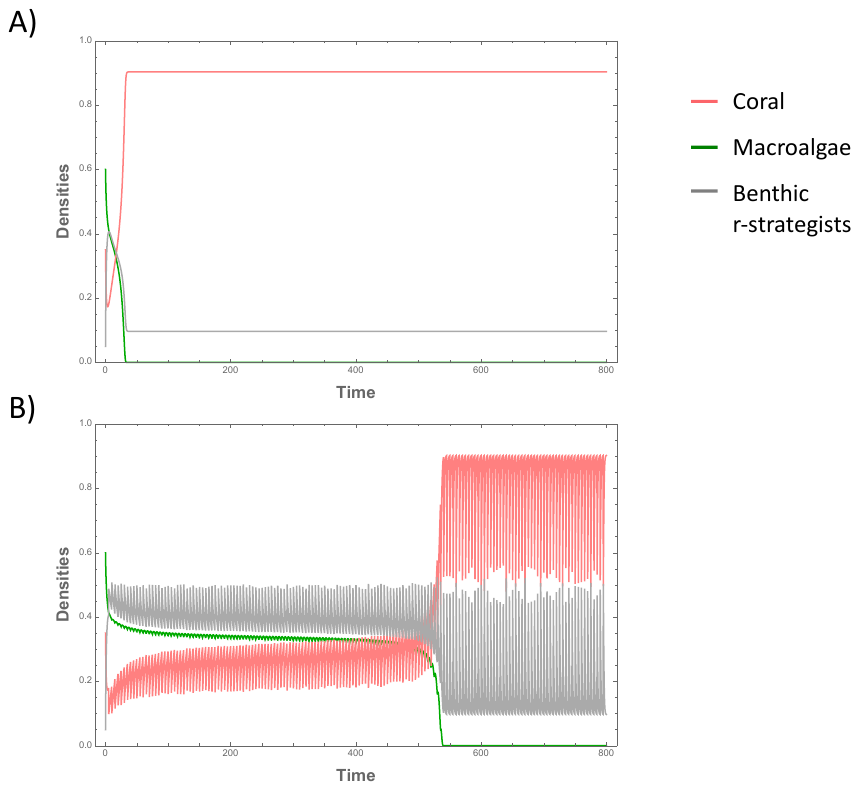

**Supplementary Figure S4.** Long transients with frequent perturbations. a) Time series without perturbations, and b) time series with perturbations. Pink line represents coral cover, green represents macroalgae, and grey is other benthic r-strategists. Parameters and initial values are constant between the two cases; *a* = 2.0, *y* = 0.7, *r* = 1.8, *m* = 0.15, *N*_0_ = 0.5, *c* = 0.25, g=0.41, Nt=0.55, and perturbation size in (b) is -0.47% every 5 time units.

These disturbances can also directly lead to climate-driven coral collapse by pushing the system out of the basin of attraction for the coral equilibrium. As disturbance frequency and/or magnitude increases the likelihood of climate-driven coral collapse also increases, meaning a bistable configuration in a deterministic scenario is much more likely to result in an algae-dominated state regardless of initial conditions. Supplementary Figure C5 shows an example of climate-driven coral collapse under a bistable configuration.

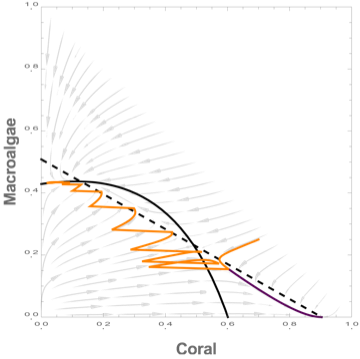

**Supplementary Figure S5.** Climate-driven coral collapse – two trajectories (with and without noise, orange and purple, respectively) with the same initial values and differential outcomes. The red trajectory shows that noise alone can cause climate-driven coral collapse even when starting within the basin of attraction to the coral-only (M=0) stable equilibrium. Here, *a* = 2.3, *y* = 0.7, *r* = 1.8, *m* = 0.15, *N*_0_ = 0.5, *c* = 0.25, g=0.4, Nt=0.55.

The result discussed in the main text regarding the presence of disturbance-driven bistability is clearly dependent on the frequency and magnitude of perturbations. Supplementary Figure C6 shows how the region of bistability (based on diverging asymptotic behaviour) changes with increasing disturbance frequency. Specifically, the upper boundary of the region of bistability increases as we increase disturbance frequency (i.e., we get bistability with higher grazing rates and lower nutrients). This region that would not be considered bistable based on the deterministic skeleton, is what we refer to as a region of climate-induced uncertainty. Also note in Supplementary Figure C6 that the lower-boundary also shifts, and this is a result of climate-driven coral collapse such that areas of parameter space that would lead to bistability in a deterministic scenario only lead to algae-dominated outcomes.

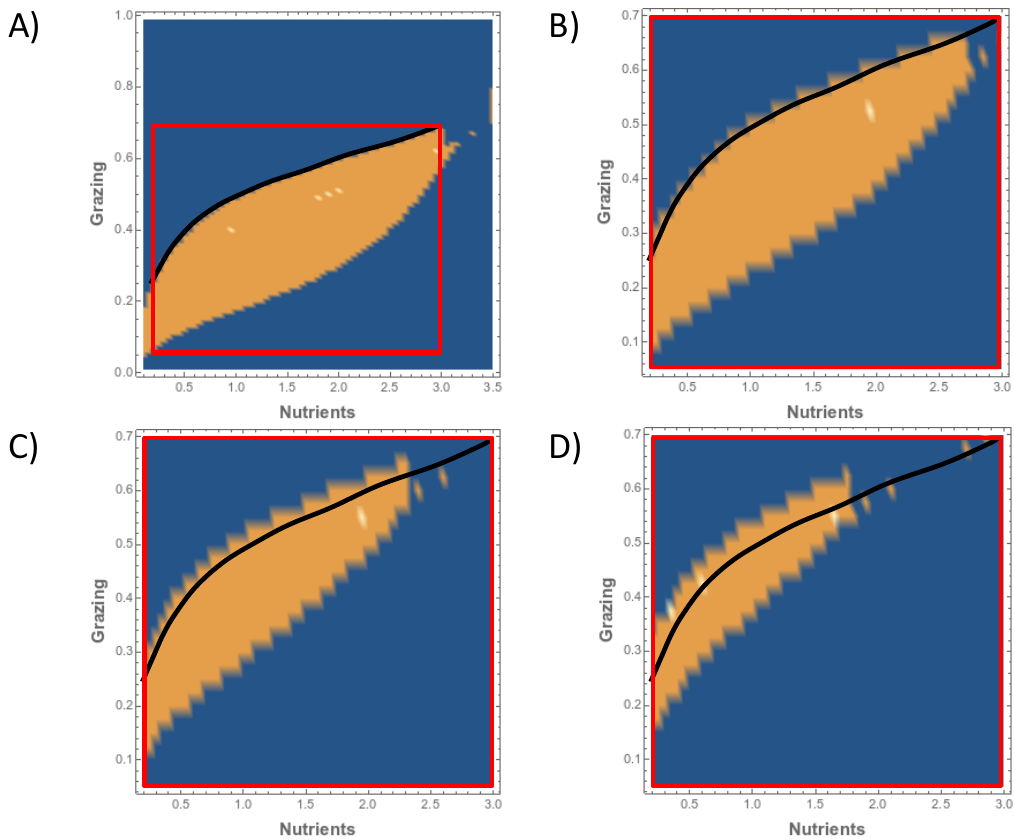

**Supplementary Figure S6.** Disturbance frequency and presence of disturbance-driven bistability. Blue areas of parameter space indicate only one asymptotic solution (either coral or macroalgae dominated) and yellow represents areas of bistability. A) disturbance (-50% coral) every 10 time units. Note that this is only marginally different from the deterministic equilibrium structure. The parameter region outlined in red indicates the area of interest shown in panels B-D, and the black line traces out the upper boundary of the bistable region from (A) for comparison in B-D. B) Disturbances every 5 time units; C) Disturbances every 3 time units; and D) Disturbances every 2 time units. In all cases, *a* = 2.0, *y* = 0.7, *r* = 1.8, *m* = 0.15, *N*_0_ = 0.5, *c* = 0.25.

**S.3 Case study from Barbadian west coast reefs**

Data for benthic cover was collected from various sources, as described in Supplementary Table S1 and in the main text.

***Nutrient measurements***

Tomascik and Sander (1985) took various nutrient measurements along the west coast of Barbados in 1982/83. These included NO_2_, NO_3_, and NH_4_, PO_4_, and Chlorophyll a. Other researchers, including Allard (1994) did not report NH_4_ measurements, so we were unable to include a comprehensive estimate of total dissolved inorganic nitrogen (DIN) for the 1992/93 studies. However, using Tomascik and Sander’s data, we calculated that the average contribution of NH_4_ to total DIN was 50.5% (standard deviation 7.5%). Thus, for a comparable estimate of DIN over time, we doubled the NO_3_ measurements reported by Vezina (1974) (later reported in Tomascik and Sander (1985)), and Allard (1994) at each sampled location along the west coast of Barbados. Below, we report all measurements available from these sources from 1972-1993, along with known nutrient thresholds for healthy coral reefs (Lapointe 1997, Bell et al. 2014, Lapointe et al. 2019) (Supplementary Figure S7). We can see that various nutrients are consistently above thresholds. Note that nitrogen is near or above the suggested total DIN threshold even without accounting for missing NH_4_ measurements.

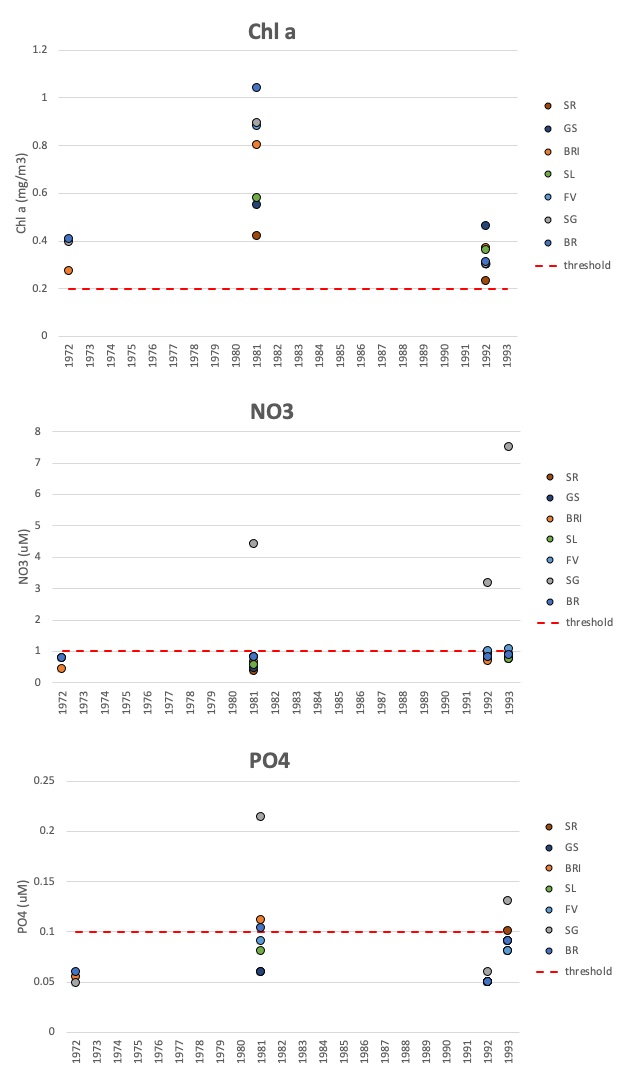

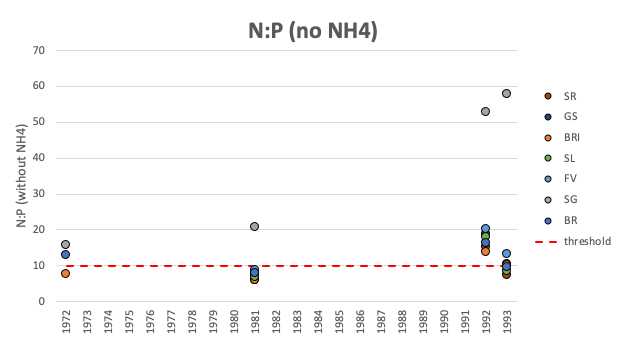

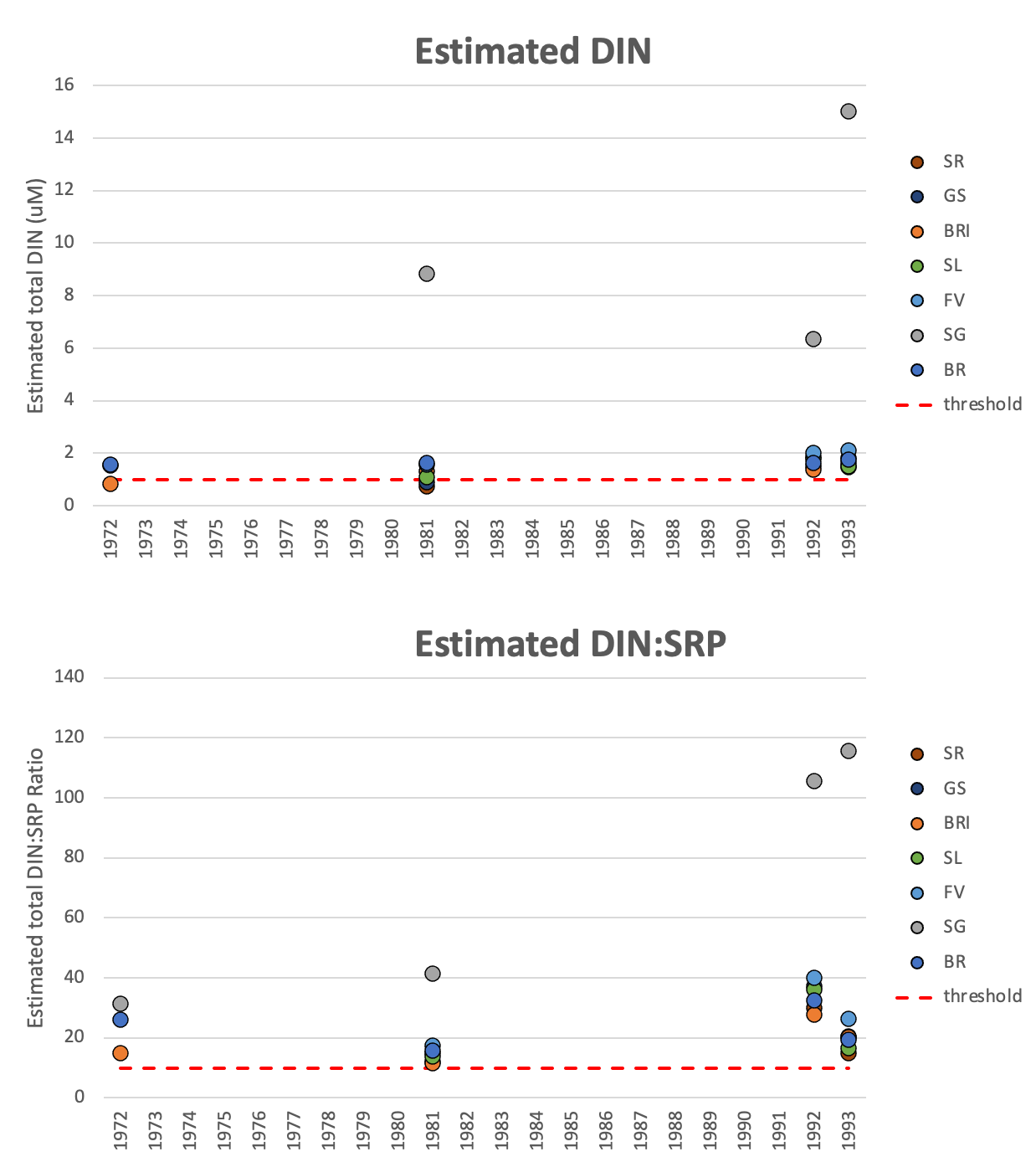

**Supplementary Figure S7.** Changes in various nutrient measurements along the west coast of Barbados. From North to South: SR = Sandridge, GS = Greensleeves, BRI = Bellairs Research Institute (South Bellairs Reef), SL = Sandy Lane, FV = Fitts Village, SG = Spring Gardens, BR= Brighton. Data from Vezina(Vezina 1974), Tomascik and Sander(Tomascik and Sander 1985), and Allard(Allard 1994), and thresholds based on Bell et al.(Bell et al. 2014), Lapointe(Lapointe 1997), and Lapointe et al.(Lapointe et al. 2019). Estimated DIN values calculated as described above, for sites with repeated measurements between 1972-1993: BR, SG, and BRI.

**Supplementary Table S1.** Data sources for changes in benthic cover and nutrients along the west coast of Barbados.

| **Data** | **Reference** |
| --- | --- |
| Baseline benthic cover | Observational late-1950s baseline information from Lewis(Lewis, John 1960); 1972 data from Stearn et al.(Stearn et al. 1977); 1981 data from Mah and Stearn(Mah and Stearn 1986) |
| Coast-wide benthic cover (along gradient) | 1982/83 data from Tomascik and Sander(Tomascik and Sander 1985, 1987a, 1987b); 1992/93 data from Allard(Allard 1994) |
| Nutrients | Vezina(Vezina 1974); Tomascik and Sander(Tomascik and Sander 1985); Allard(Allard 1994) |
| Nutrient thresholds | Bell et al.(Bell et al. 2014); Lapointe(Lapointe 1997); Lapointe et al.(Lapointe et al. 2019) |

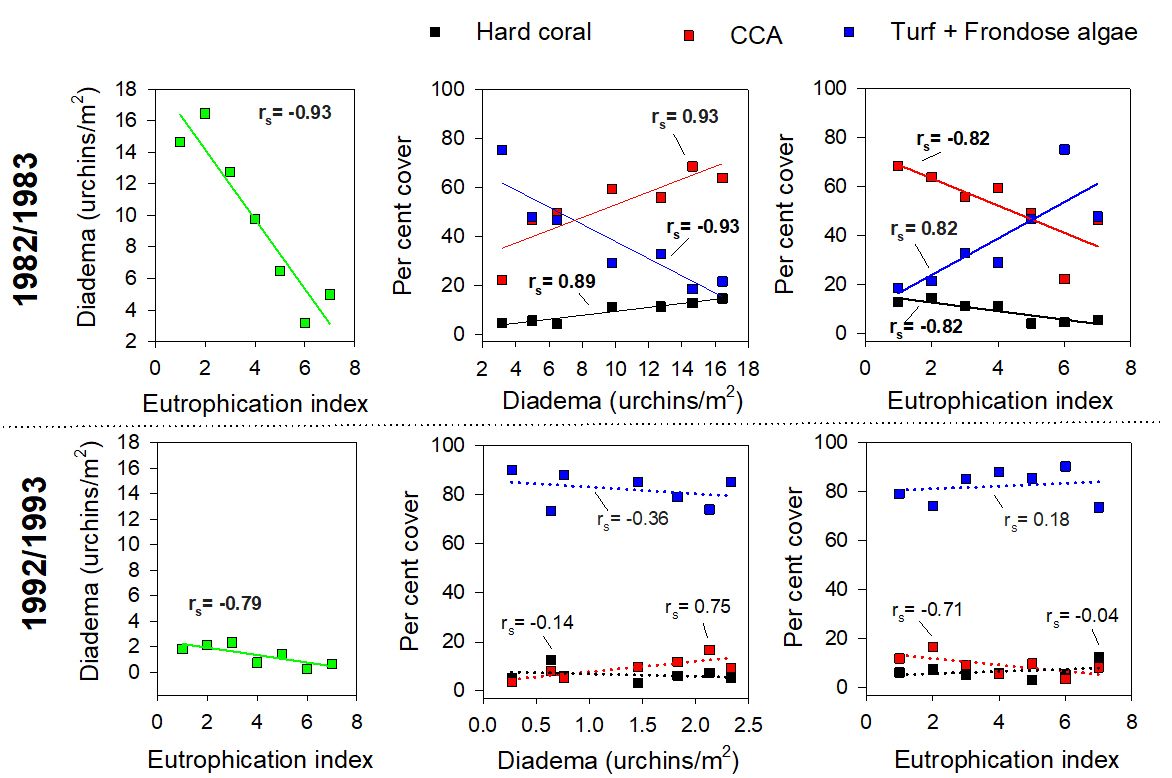
**Supplementary Figure S8.** Multi-panels showing spatial correlations across seven Barbadian reefs showing a gradient of decreasing *Diadema* density and increasing eutrophication (north-south gradient), as well as correlations between *Diadema* density and three benthic variables (hard coral cover, CCA and turf + frondose algae), and between reef eutrophication rank and the three aforementioned benthic variables. Data are shown for the pre-*Diadema* die-off period (1982/1983) and for the post-*Diadema* die off period (1992-1993). Significant Spearman rank correlations (p<0.05) are identified with bold font and solid trend lines.

**S.4 Supplementary References**

Allard, P. 1994. Changes in Coral Community Structure in Barbados: Effects of Eutrophication and Reduced Grazing Pressure. McGill University.

Bell, P. R. F., I. Elmetri, and B. E. Lapointe. 2014. Evidence of Large-Scale Chronic Eutrophication in the Great Barrier Reef: Quantification of Chlorophyll a Thresholds for Sustaining Coral Reef Communities. AMBIO 43:361–376.

Lapointe, B. E. 1997. Nutrient thresholds for bottom-up control of macroalgal blooms on coral reefs in Jamaica and southeast Florida. Limnology and Oceanography 42:1119–1131.

Lapointe, B. E., R. A. Brewton, L. W. Herren, J. W. Porter, and C. Hu. 2019. Nitrogen enrichment, altered stoichiometry, and coral reef decline at Looe Key, Florida Keys, USA: a 3-decade study. Marine Biology 166:1–31.

Lewis, John, B. 1960. The coral reefs and coral communities of Barbados, W.I. Canadian Journal of Zoology 38:1133–1152.

Mah, A. J., and C. W. Stearn. 1986. The effect of Hurricane Allen on the Bellairs fringing reef, Barbados. Coral Reefs 4:169–176.

Mumby, P. J., A. Hastings, and H. J. Edwards. 2007. Thresholds and the resilience of Caribbean coral reefs. Nature 450:98–101.

Stearn, C. W., T. P. Scoffin, and W. Martindale. 1977. Calcium carbonate budget of a fringing reef on the west coast of Barbados Part I- zonation and productivity. Bulletin of Marine Science 27:479–510.

Tomascik, T., and F. Sander. 1985. Effects of eutrophication on reef-building corals I. Growth rate of the reef-building coral Montastrea annularis. Marine Biology 87.

Tomascik, T., and F. Sander. 1987a. Effects of eutrophication on reef-building corals - II. Structure of scleractinian coral communities on fringing reefs, Barbados, West Indies. Marine Biology 94:53–75.

Tomascik, T., and F. Sander. 1987b. Effects of eutrophication on reef-building corals - III. Reproduction of the reef-building coral Porites porites. Marine Biology 94:77–94.

Vezina, R. 1974. Seawater quality and phytoplankton of inshore waters of Barbados: a study of the effects of organic pollution in a tropical environment. McGill University.
